# Aurora-A phosphorylates splicing factors and regulates alternative splicing

**DOI:** 10.1101/2020.11.04.368498

**Authors:** Arun Prasath Damodaran, Olivia Gavard, Jean-Philippe Gagné, Malgorzata Ewa Rogalska, Estefania Mancini, Thibault Courtheoux, Justine Cailloce, Agnès Mereau, Guy G. Poirier, Juan Valcárcel, Erwan Watrin, Claude Prigent

## Abstract

Aurora-A kinase is well known to regulate progression through mitosis. However, the kinase also performs additional functions that could explain the failure of its inhibitors to be effective in cancer treatments. To identify these functions, we applied a proteomics approach to search for interactors of Aurora-A. We found a large number of proteins involved in pre-mRNA splicing, strongly suggesting an important role for Aurora-A in this biological process. Consistently, we first report the subcellular localization of Aurora-A in nuclear speckles, the storehouse of splicing proteins. We also demonstrate direct interaction of Aurora-A with RRM domain-containing splicing factors such as hnRNP and SR proteins and their phosphorylation *in vitro*. Further, RNA-sequencing analysis following pharmacological inhibition of Aurora-A resulted in alternative splicing changes corresponding to 505 genes, including genes with functions regulated by Aurora-A kinase. Finally, we report enrichment of RNA motifs within the alternatively spliced regions affected by Aurora-A kinase inhibition which are bound by Aurora-A interacting splicing factors, suggesting that Aurora-A regulates alternative splicing by modulating the activity of these interacting splicing factors. Overall our work identified Aurora-A as a novel splicing kinase and for the first time, describes a broad role of Aurora-A in regulating alternative splicing.

## INTRODUCTION

Aurora-A belongs to the family of serine/threonine kinases and is one among three Aurora kinases (-A, -B and -C) in mammals (1). The first member of the family was identified in yeast, that possess only one Aurora encoded by the gene Ipl1 (for Increase in ploidy 1) that was demonstrated to be involved in the regulation of ploidy (2, 3). Aurora-A was later identified as a protein kinase that localizes to centrosome, is overexpressed in cancers, and is required for bipolar spindle assembly during mitosis (4–6). The level of Aurora-A remains very low during G1 phase and starts to increase in S phase to reach its maximum protein levels at mitosis (7, 8). Aurora-A regulates, not only mitotic spindle assembly, but multiple cell cycle events such as centrosome maturation, mitotic entry, cytokinesis and mitotic exit and, thereby, enables proper cell division in cooperation with additional cell cycle-regulating proteins (9). Aurora-A is also an oncogene that induces tumorigenesis by inducing a long period of genetic instability (10). While its role in cell cycle control through phosphorylation of mitotic proteins is well established, recent studies have unveiled novel roles beyond cell cycle regulation such as mitochondrial dynamics, gene regulation and translation (11–13).

Aurora-A is over-expressed in multiple epithelial cancer types such as breast, ovarian, gastrointestinal, colorectal, prostate and lung cancers (14, 15). Although normal cells must tightly regulate the level of Aurora-A to ensure normal mitosis, cancer cells seem to find an advantage in overexpressing it, without compromising their cell division too much. This overexpression might rather help in cancer development by promoting cell proliferation, cell survival, stemness, metastasis and preventing apoptosis (8). The immediate realization of the role and importance of Aurora-A in tumour development and progression has made it a prime target for cancer treatment. Though many drugs inhibiting Aurora-A kinase have been identified, none of them has cleared the phase III clinical trials due to their toxicity (16). Therefore, it appears essential to better understand all the functions of Aurora-A in order to identify new ways to target the kinase and to avoid toxicity. In an attempt to gain insights into the full spectrum of Aurora-A functions, we performed mass spectrometry-based analysis of Aurora-A interactome, which revealed several splicing proteins as interacting partners of Aurora-A.

RNA splicing involves the removal of introns and ligation of flanking exons to form mature mRNA (17, 18). Pre-mRNA splicing is carried out by a complex and dynamic machinery called spliceosome, which assembles on a pre-mRNA in a stepwise manner forming multiple intermediate spliceosomal complexes (19, 20). To date, more than 200 proteins are known to be associated with the spliceosome at different steps of the splicing process (21–23). 141 highly abundant splicing proteins were categorized as core-components of the spliceosome and include the snRNPs (small nuclear ribonucleoprotein). Conversely, 103 less abundant proteins were categorized as non-core proteins and include mostly regulatory proteins such as SR (Serine and arginine-rich) proteins and hnRNPs (Heterogeneous nuclear ribonucleoproteins) (24, 25). In the present article, we report the identification of splicing factors as novel interactors of Aurora-A and, further, provide evidence that Aurora-A kinase activity is required for regulating alternative splicing, likely through modulation of the activity of its interacting factors, altogether describing Aurora-A as a novel splicing kinase.

## MATERIALS AND METHODS

### Expression vectors and molecular cloning

The plasmid constructs were generated using Gibson Assembly Master Mix (New England Biolabs). The cDNAs of splicing proteins were PCR amplified and was subcloned into pGEX4T1 vector. All plasmid constructs were verified by sequencing. All the 12 pCS2+ FLAG-tagged SRSF constructs for expression in mammalian cells were a kind gift from Dr. Wei Wu (26). All the GST tagged SRSF constructs for expression in bacterial cells were a kind gift from Dr. Evelyne Manet (27). The domain deletion mutants of pCS2+ FLAG-tagged SRSF1 construct were generated using Q5^®^ Site-Directed Mutagenesis Kit (New England Biolabs). The primers sequences are given in **Supplementary Table S1**.

### Cell culture and transfection

U2OS, HEK293 and HeLa cells were grown in Dulbecco’s modified Eagle’s medium (DMEM)-GlutaMAX™ -I (Sigma-Aldrich) supplemented with 10% fetal bovine serum (GE Healthcare) and 1% penicillin–streptomycin (GE Healthcare). For transfection experiments, cells were grown until they reached 60% confluency and the plasmids were transfected using Lipofectamine-3000 (ThermoFisher scientific) transfection reagents according to the manufacturer’s instructions.

### Cell synchronization and inhibition of Aurora-A

For synchronization of HeLa cells at G1 phase, we used a double thymidine block method where Aurora-A inhibitor (MLN8237) was added immediately after the first thymidine release when the cells were in G1 phase. Then, non-mitotic cells were collected after 24 hours of Aurora-A inhibitor (50 nM) treatment. For synchronization at G2 phase, we released cells for 6 hours from double thymidine block. Aurora-A inhibitor was added 6 hours after the first thymidine release when the cells were in G2 phase. Then, non-mitotic cells were collected after 24 hours of Aurora-A inhibitor treatment. For synchronization at mitotic phase, we released cells for 12 hours from double thymidine block. Aurora-A inhibitor was added 12 hours after the first thymidine release when the cells were in late G2 or mitotic phase. Then, mitotic cells were collected after 24 hours of Aurora-A inhibition. Nocodazole (3.3 μM) was added two hours prior to harvesting cells to collect a maximum number of mitotic cells.

### Generation of U2OS-GFP-Aurora-A with shRNA-Aurora-A stable cell line

Stable depletion of endogenous Aurora-A in U2OS-GFP-Aurora-A cell line (28) was obtained by the transfection of shRNA directed against the 3’ UTR region of Aurora-A (clone NM_003600.x-985s1c1, Sigma) followed by clonal dilution and selection of Aurora-A depleted cells with 2 μg/ml of puromycin.

### Western blot analysis

The cells were washed with PBS twice and scraped directly in 2X Laemmli sample buffer and boiled at 95°C for 5 minutes. The protein concentration was quantified using the Protein Quantification Assay kit (Macherey-Nagel). The protein samples were resolved using 4–15% Mini-PROTEAN™ TGX Stain-Free™ Protein Gels (Bio-Rad) and transferred onto PVDF membrane (GE Healthcare) and analysed by western blotting. The following primary antibodies were used: rat anti-a-tubulin (MAB1864, Millipore, 1:10000), mouse anti-Aurora-A (clone 5C3, 1:100 (29)), rabbit anti-phospho-Aurora-A/B/C (clone D13A11, 2914, Cell Signaling Technology, 1:500), mouse anti-ASF/SF2 (clone 96, 32-4500, Thermo Fisher Scientific, 1:10000), mouse anti-GST tag (G1160, Sigma 1:2000), mouse anti-His tag (HIS.H8 / EH158, Covalab, 1:2500), mouse anti-FLAG tag (clone M2-F1804, Sigma, 1:1000), mouse anti-GFP (clones 7.1 and 13.1, Roche Applied Science, Indianapolis, IN, 1:1000). Corresponding secondary horseradish peroxidase-conjugated antibodies were used: anti-mouse and anti-rabbit (Jackson Immuno Research Laboratories) and anti-rat (A110-105P, Bethyl Laboratories). Membranes were incubated with a lab-made enhanced chemiluminescence reagent containing 100 mM Tris (pH 8.5), 13 mg/ml coumaric acid (Sigma), 44 mg/ml luminol (Sigma) and 3% hydrogen peroxide (Sigma). Chemiluminescence signals were captured on a film (CP-BU new, Agfa Healthcare) with a CURIX 60 developer (Agfa Healthcare).

### Affinity-purification of Aurora-A-interacting proteins

Multiple biological and technical replicates were used to generate a highly specific dataset of Aurora-A-associated proteins. Experiments were performed in triplicates and injected in two fractions for LC-MS/MS analysis (a total of 6 replicates for control experiments and 12 replicates corresponding to GFP-Aurora-A affinity-purification extracts). In order to assess the non-specific binding to our affinity-purification matrix (i.e. Dynabeads coupled to anti-GFP antibodies), whole cell extracts of wild-type human osteosarcoma U2OS cells were used. For affinity-purification of Aurora-A-associated protein complexes, whole cell extracts of U2OS cells stably expressing a GFP-tagged version of human Aurora-A were used. Wild-type and GFP-Aurora-A expressing U2OS cells were seeded onto three 150 mm cell culture dishes and grown up to 75% confluency. Cells were incubated with 1 μg/ml paclitaxel for 18 h to cause a mitotic arrest. Two PBS washes were carried out prior to harvesting the cells with disposable cells scraper. All the affinity-purification steps were carried out at 4°C unless mentioned otherwise. Three millilitres/plate of lysis buffer (50 mM HEPES pH 7.5, 150 mM NaCl, 0.3% CHAPS and supplemented with cOmplete™ EDTA-free protease inhibitor cocktail (Roche) and 100 units mL Benzonase^®^ nuclease (Sigma-Millipore)) were used during cells scraping. Cells were kept on ice and lysed for 30 min on a rotating device. Cell extracts were centrifuged for 5 min at 10,000 rpm to remove cellular debris. Immunoprecipitation experiments were performed using Dynabeads™ magnetic beads covalently coupled with Protein G (Life Technologies). The Dynabeads™ were washed once with 2 ml of PBS and coated with 15 μg of mouse monoclonal anti-GFP antibody (Roche Applied Science). The beads were finally washed three times with 2 ml of lysis buffer and added to the whole cell extract for a 2 h incubation with gentle rotation. Samples were then washed three times with 1 volume of lysis buffer for 10 min. Beads were resuspended in a 75 mM ammonium bicarbonate solution (pH 8.0) and protein complexes were directly digested on-beads by the addition of 2 μg of a Trypsin/Lys-C mixture (Promega). Peptides were isolated on C18 resin tips according to the manufacturer’s instructions (Thermo Fisher Scientific) and dried to completion in a speed vac evaporator.

### Liquid-chromatography tandem mass spectrometry (LC-MS/MS)

All samples were processed on an Orbitrap™ Fusion™ Tribrid™ mass spectrometer (Thermo Fisher Scientific). Peptide samples were separated by online reversed-phase (RP) nanoscale capillary liquid chromatography (nanoLC) and analysed by electrospray mass spectrometry (ESI MS/MS). The experiments were performed with a Dionex UltiMate 3000 nanoRSLC chromatography system (Thermo Fisher Scientific / Dionex Softron GmbH, Germering, Germany) connected to the mass spectrometer equipped with a nanoelectrospray ion source. Peptides were trapped at 20 μl/min in loading solvent (2% acetonitrile, 0.05% TFA) on a 5 mm x 300 μm C18 pepmap cartridge pre-column (Thermo Fisher Scientific / Dionex Softron GmbH, Germering, Germany) for 5 minutes. Then, the pre-column was switched online with a self-made 50 cm x 75μm internal diameter separation column packed with ReproSil-Pur C18-AQ 3 μm resin (Dr. Maisch HPLC GmbH, Ammerbuch-Entringen, Germany) and the peptides were eluted with a linear gradient from 5-40% solvent B (A: 0,1% formic acid, B: 80% acetonitrile, 0.1% formic acid) in 60 minutes, at 300 nL/min. Mass spectra were acquired using a data dependent acquisition mode using Thermo XCalibur software version 3.0.63. Full scan mass spectra (350 to 1800m/z) were acquired in the orbitrap using an AGC target of 4e5, a maximum injection time of 50 ms and a resolution of 120 000. Internal calibration using lock mass on the m/z 445.12003 siloxane ion was used. Each MS scan was followed by the acquisition of fragmentation spectra of the most intense ions for a total cycle time of 3 seconds (top speed mode). The selected ions were isolated using the quadrupole analyser in a window of 1.6 m/z and fragmented by Higher energy Collision-induced Dissociation (HCD) with 35% of collision energy. The resulting fragments were detected by the linear ion trap in rapid scan rate with an AGC target of 1e4 and a maximum injection time of 50 ms. Dynamic exclusion of previously fragmented peptides was set for a period of 20 sec and a tolerance of 10 ppm.

### Data processing

Mass spectra data generated by the Orbitrap™ Fusion™ Tribrid™ instrument (*.raw files) were analysed by MaxQuant (version 1.5.2.8) using default settings for an Orbitrap instrument. Andromeda was used to search the MS/MS data against FASTA-formatted Homo sapiens Uniprot Reference Proteome UP000005640 (70 611 entries) complemented with GFP-Aurora-A amino acid sequence and a list of common contaminants maintained by MaxQuant and concatenated with the reversed version of all sequences (decoy mode). The minimum peptide length was set to 7 amino acids and trypsin was specified as the protease allowing up to two missed cleavages. The false discovery rate (FDR) for peptide spectrum matches (PSMs) was set at 0.01. The following mass additions were used as variable modifications: oxidation of methionine [+15.99491 Da], deamidation of asparagine and glutamine [+0.98401 Da], N-terminal acetylation [+42.01056 Da] and phosphorylation of serine and threonine [+97.97689 Da]. MaxQuant output files were imported in Scaffold (version 4.6.1, Proteome Software Inc) to validate MS/MS based peptide and protein identifications. Peptide and protein identifications were accepted if they could be established at greater than 95% probability to achieve an FDR less than 1%. Common contaminants (i.e. keratins, serum albumin, IgGs, trypsin, ribosomal subunit proteins, etc) were removed from the output file. Proteins found as non-specific binders of magnetic beads coupled to anti-GFP antibodies were removed from the GFP-Aurora-A-specific dataset unless they are assigned with 10 times more peptide spectrum matches (PSMs).

Finally stringent cut-off was applied to obtain the specific interactors of Aurora-A where the proteins with sequence coverage less than 5%, Label free quantification (LFQ) for control not equal to 0 and LFQ for Aurora-A equal to 0 were filtered out.

### Gene ontology enrichment analysis

The Gene Ontology enrichment analysis was carried out using the tool g:Profiler (30) using the GO biological processes and following parameters (Significance threshold - Benjamini and Hochberg method; User threshold: 0.05; minimum term size is 5 and maximum term size is 350). The functional GO terms were visualized using Enrichment Map plugin v3.3.0 (31) available in the Cytoscape v3.8.0 (32) with following parameters (FDR q-value cutoff of 0.01; p-value cutoff of 0.001; metric: overlap; metric cutoff: 0.8). The functional GO terms were arranged (mcl clustering) based on similarity using AutoAnnotate plugin v1.3.3 (33) available in the Cytoscape v3.8.0. In enrichment maps, node size is proportional to the gene-set size, border width colour is proportional to FDR (q-value) and edge width is proportional to the number of genes shared between GO categories. Alternately, gene ontology enrichment analysis was carried out using the DAVID tool (34, 35). For all GO enrichment analysis, the default background (human proteome) as assigned by the tools was used.

### Construction of protein-protein interaction network

The protein-protein network was constructed using Cytoscape v3.8.0 and the integrated network query using STRING plugin v1.5.1 (36) with the Confidence (score) cutoff of 0.9 and maximum additional interactors of 0. The sub-networks or clusters were visualized using mcode plugin v1.6.1 (37) available with Cytoscape v3.8.0.

### Expression and purification of recombinant proteins

E.coli strain BL21 (DE3) was transformed with GST fusion plasmids encoding HNRNPC, RALY, PCBP2, PTBP1, RS26, eIF4A3, SRSF1, SRSF3 and SRSF7 or HIS fusion plasmid encoding Aurora-A. Cells were grown till the OD600 reached 0.6 in 250 ml LB media supplemented with antibiotics (ampicillin 100 μg/ml or kanamycin 50 μg/ml) and expression of the protein was induced for 4 hours with 1 mM isopropyl 1-thio-B-D-galactopyranoside (Sigma) at 18°C. Cells were collected and sedimented by centrifugation for 20 min at 4000 rpm (4°C). The cell pellet for HIS-tagged protein purification was resuspended in 10 ml of Buffer A (20 mM Tris pH 7.5, 500 mM NaCl, 20 mM imidazole) containing 1X protease inhibitors (EDTA free protease inhibitor cocktail, Roche). Whereas for purification of GST tagged proteins, cell pellets were resuspended in 10 ml of Buffer B (0.1% Triton X100 in PBS) supplemented with 1X protease inhibitors. The cells were then sonicated at 50% amplitude for 3 min (30 sec pulses) and cell lysates were centrifuged at 10000 rpm for 30 min at 4°C. The supernatant was incubated with TALON His-tag purification resin (TaKaRa) or glutathione agarose beads (Sigma-Aldrich) for 30 min in a rocker at 4°C. Beads were then washed thrice with buffer and eluted with 250 mM imidazole (for HIS-tagged proteins) or 5 mM reduced glutathione (for GST-tagged proteins). Protein profiles from the eluted fractions were estimated by SDS-PAGE followed by Coomassie staining.

### *In vitro* interaction assay

For GST pull-down assays, the GST-tagged hnRNP and HIS-tagged Aurora-A were incubated in interaction buffer (containing 50 mM Tris-HCl, pH 7.5, 100 mM KCl, 10 mM MgCl_2_, 5% glycerol, 1 mM DTT and 0.25% NP40) for 2 hours in a rocker at 4°C and the complex was pulled down using Glutathione Sepharose 4B beads. The interaction of splicing proteins with Aurora-A was analysed by western blots against anti-GST and anti-HIS or anti-Aurora-A antibodies.

For FLAG pull-down assays, FLAG-tagged SR proteins (SRSF) were expressed in HEK293 cells and the cells were lysed with M2 lysis buffer (50 mM Tris-base (pH 7.4), 150 mM NaCl, 10% glycerol, 1% Triton X-100 and 1 mM EDTA). The FLAG-tagged SR proteins were incubated with FLAG-M2 beads and were washed thrice with M2 lysis buffer. The bound FLAG-tagged SR proteins were incubated with recombinant HIS-tagged Aurora-A for 2 hours in a rocker at 4°C. The beads containing protein complexes were again washed thrice and eluted with 100 μg/ml of FLAG peptide. The interaction of SR proteins with Aurora-A was analysed by western blots against anti-FLAG and anti-HIS or anti-Aurora-A antibodies.

### *In vitro* kinase assay

Recombinant GST-tagged splicing proteins and HIS-tagged Aurora-A were incubated in kinase buffer (containing 50 mM Tris-HCl, pH 7.5, 50 mM NaCl, 1 mM DTT, 10 mM MgCl_2_ and 100 μM ATP) in presence of [γ-^32^P]-ATP at 37°C for 30 minutes. The reaction was stopped by the addition of 2X Laemmli sample buffer and boiled at 95°C for 5 min. Phosphorylated splicing proteins were resolved by SDS-PAGE on 10% gels and visualized by autoradiography.

### Immunofluorescence staining and microscopy

Cells grown on coverslips were fixed with methanol (−20°C) for 5 minutes. The cells were then blocked with 5% FBS in PBS and incubated with SC35 primary antibody (S4045, Sigma) at 1:2000 dilution in blocking solution for 1 hour at room temperature. Cells were washed thrice in PBS containing 0.05% Tween 20 and incubated with secondary antibody (anti-mouse antibody conjugated to Alexa 647 or Alexa 546, Thermo Fisher Scientific) at 1:4000 dilution and GFP nano-booster (Chromotek) at 1:1000 dilution in blocking solution for one hour at room temperature. The cells were then washed and the DNA was stained using Hoechst 33342 (Thermo Fisher Scientific) or DAPI. Finally, the coverslips were mounted using ProLong^®^ Gold antifade reagent (Thermo Fisher Scientific). High-resolution images were taken with the LSM800 Airyscan confocal microscope (Zeiss Inc.) equipped with 63X/1.4 Plan Apochromat Oil DIC objective and Airyscan detector (airyscan mode). The images were processed with FIJI (ImageJ) software.

For live cell imaging, stable U2OS cells expressing GFP-Aurora-A and shRNA-Aurora-A were grown in 35 mm glass-bottomed dishes (Matek Corporation) Z-sweeps of 0.3 μm sections were acquired for both GFP and DAPI channels. Subsequent analysis was carried out using Fiji (ImageJ) software.

### Flow cytometry analysis and mitotic index calculation

After the cell cycle specific synchronization, HeLa cells were washed twice with PBS and was resuspended in 70% ice-cold methanol and incubated at 4°C for 1 hour. Pellets were washed twice with PBS and resuspended in PI buffer containing 50 μg/ml propidium iodide, 10 mM Tris pH7.5, 5 mM MgCl_2_, 200 μg/ml RNase A and incubated at 37°C for 30 minutes. The percentage of cells in each cell cycle phase was calculated with a Fortessa X-20 LSR (Becton Dickinson). The mitotic index of FACS samples was calculated in order to discriminate the cells in G2 and mitotic phase. The samples prepared for FACS analysis were mounted on a glass slide and counted using a fluorescence microscope (DMRXA2 Leica). At least 100 cells were counted for each condition.

### RNA-sequencing analysis

Total RNA was isolated from two replicates of the synchronized HeLa cells using the RNeasy Plus Mini Kit (QIAGEN) as per the manufacturer’s protocol. While isolating total RNA for RNA-seq analysis, on-column digestion of DNA was additionally performed using RNase-Free DNase Set (QIAGEN) as a precautionary measure. The mRNA-seq libraries were sequenced using 125bp paired end reads on Illumina HiSeq 2500 lanes. The Library preparation and sequencing were performed in the CRG Genomics Unit. The quality of the RNA sequencing data was evaluated using the MultiQC tool. The reads were aligned using the Star alignment program to the hg38 human reference genome. The read counts were corrected using edgeR upper quartile normalization factors. Differential gene expression analysis between Aurora-A inhibited and control conditions were carried out using edgeR tool (38) and a final cut-off for log_2_-fold change of 0.5 and adjusted P-value of 0.05 were applied. Alternative splicing analysis was carried out using *vast-tools* v2.2.0 pipeline (39, 40) which is available in github (https://github.com/vastgroup/vast-tools). PSI values for two replicates were quantified for all types of events (cassette exons, alternative 3’ and 5’ splice sites and retained introns). A minimum ΔPSI of 15% was required to define differentially spliced events with a minimum range of 5% between the PSI values of the two samples (control and Aurora-A inhibited samples). The RNA motif enrichment for splicing factors/RBP were carried out using *Matt-tool* v1.3.0 (41) with the differentially spliced events (|ΔPSI| > 15) and unregulated splicing events as inputs for computing the enrichment profile.

## RESULTS

### Shotgun proteomics analysis of Aurora-A interaction network reveals an expanded suite of potential binding partners and substrates

To identify novel interacting partners of Aurora-A, we performed a comprehensive analysis of proteins found in Aurora-A affinity-purifications (**Figure 1A**). For this, we used previously established bone osteosarcoma U2OS cell line expressing a GFP-tagged version of Aurora-A under the control of its own promoter (28) to generate a distinct cell line that, in addition to expressing GFP-Aurora-A, expresses shRNA that allows RNAi-mediated silencing of endogenous Aurora-A mRNA (**Figure 1B**). In this novel cell line, decrease of endogenous Aurora-A leads to an increase of exogenous GFP-Aurora-A protein level (**Figure 1B**) as previously observed (28), possibly through compensatory mechanism. In addition, this cell line displays localisation of GFP-Aurora-A to the interphase centrosome and mitotic spindle (**Supplementary Figure S1A**) and normal cell cycle profile identical to that of wild-type U2OS cells (**Supplementary Figure S1B**) ruling out any abnormal phenotype in this cell line. Wild type U2OS cells and U2OS cells expressing GFP were used as negative controls to identify Aurora-A-specific interactors. As Aurora-A protein level reaches maximum at mitosis and in order to capture a maximum number of interacting partners all three cell lines were synchronized in mitosis using paclitaxel (taxol). Affinity-purification was performed using anti-GFP monoclonal antibody coupled to Protein-G-coated magnetic beads (**Figure 1C**). The presence of active GFP-Aurora-A in the affinity-purified sample was revealed by western blot analyses using anti-Aurora-A, anti-pT288-Aurora-A and anti-GFP antibodies (**Figure 1D**). Further, affinity-purified protein complexes were directly digested on beads and subjected to liquid chromatography coupled to tandem mass spectrometry (LC-MS/MS). The global proteomics analysis (see methods) identified 407 specific interacting protein-partners of Aurora-A (**Figure 1E**). Notably, these included 36 previously validated human interactors of Aurora-A out of 374 Aurora-A interactors retrieved from the interaction repository BioGRID (**Figure 1F and Supplementary Table S2**). These known interactors include well-characterized Aurora-A effectors such as TPX2 (42, 43), CEP192 (44), PPP6C (45), PPP1CB (46) and WDR62 (47) providing solid indications that our proteomics dataset is likely to contain other relevant interactions. Overall, large-scale shotgun proteomics analysis of Aurora-A-associated complexes generated a dataset of 371 potential, yet uncharacterized, interaction partners or substrates selectively targeted by Aurora-A (**Figure 1F and Supplementary Table S2**).

**Figure 1.**
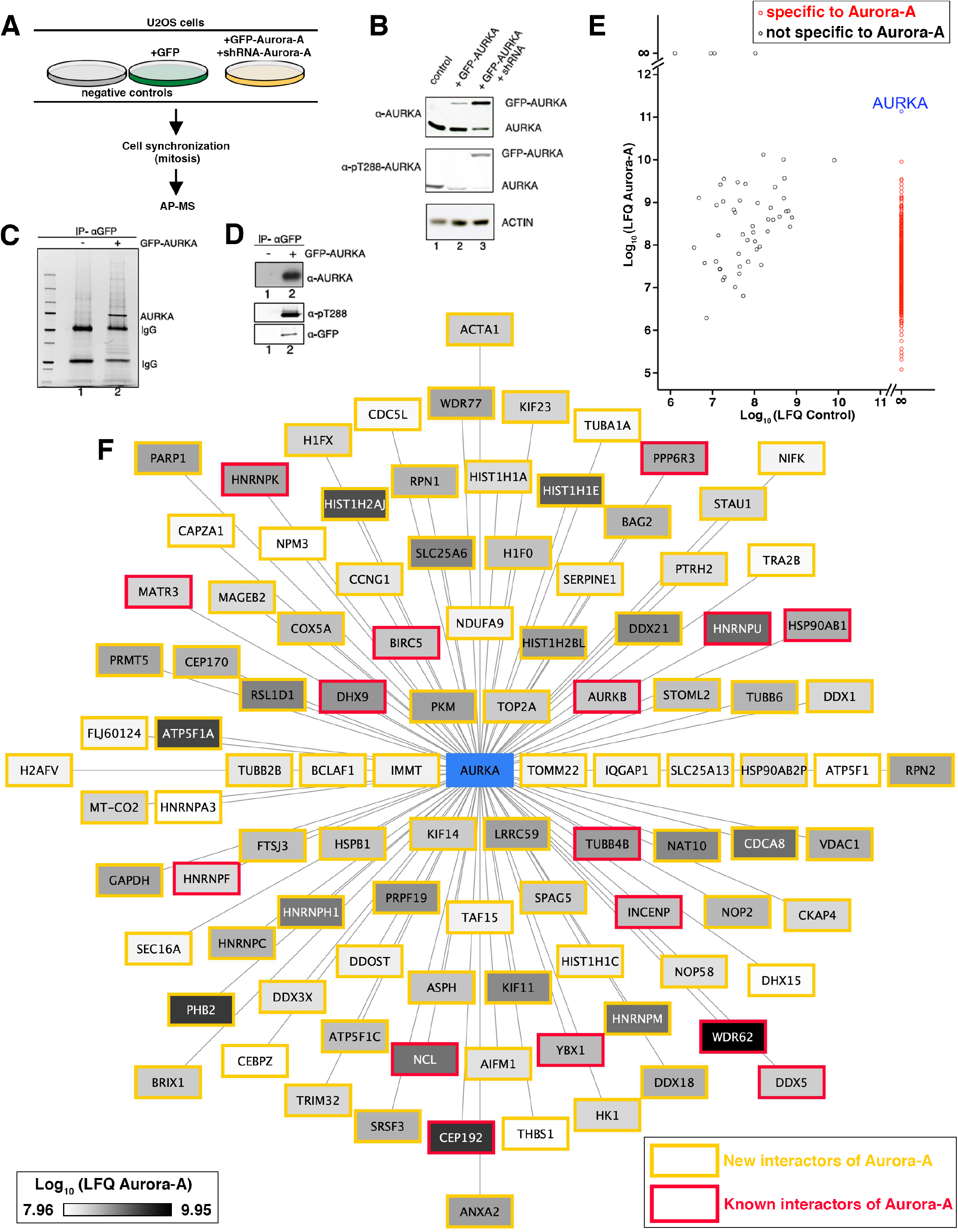
Identifying new interacting partners of Aurora-A. (**A**) Schematic representation of the strategy used to isolate and identify Aurora-A interacting proteins in human cells. Two negative controls (wild-type U2OS cells and U2OS cells expressing GFP) and U2OS with stable expression of GFP-Aurora-A and shRNA-Aurora-A were used. Cells were synchronized in mitosis by overnight incubation with Taxol. GFP and GFP-Aurora-A were isolated by affinity-purification coupled to mass spectrometry (AP-MS). (**B**) Western blot analysis of the expression level of GFP-Aurora-A. Protein extracts prepared from control U2OS cells (1), from cells expressing GFP-Aurora-A (2), and from cells expressing GFP-Aurora-A and Aurora-A-shRNA (3) were analysed by Western blot. Endogenous Aurora-A and GFP-Aurora-A protein were detected by anti-Aurora-A antibody (top panel) while the activated kinase by anti-phospho-Thr288 (middle panel). Actin was used as a loading control (lower panel). (**C**) Profiles of proteins co-pulled down by anti-GFP antibody from control U2OS cells (1) and U2OS cells expressing GFP-Aurora-A and Aurora-A-shRNA (2). The gel is showing the co-pulled down proteins stained with Sypro Ruby. (**D**) Control by western blot of affinity-purified GFP-Aurora-A with anti-Aurora-A antibody (top panel), anti-pThr288 antibody (middle panel) and anti-GFP antibody (lower panel). (**E**) The common contaminants and proteins with sequence coverage less than 5% were removed from the global proteomic dataset. This filtered dataset was then subjected to scatter plotting where x and y axes represent the logarithmized Label-free quantification (LFQ) values for proteins in control and Aurora-A affinity-purification respectively. The red circles indicate the interacting partners specific to Aurora-A. (**F**) The top 100 proteins in the filtered proteomics dataset of 407 Aurora-A specific interacting proteins as visualized in Cytoscape.

### Gene ontology analysis of Aurora-A interactome reveals enrichment for proteins related to RNA splicing

The non-cell cycle-related functions of Aurora-A are poorly understood. To gain insights on the new potential functions of Aurora-A, the interactome data of Aurora-A was subjected to Gene Ontology (GO) enrichment analysis using g:Profiler online tool. This revealed 225 highly significant and redundant biological processes (FDR<0.01) that resulted from querying a large fraction (365/407 proteins or ~90%) of the Aurora-A interactors. These redundant biological processes were grouped to closely related functions by constructing a similarity-based network using the GO enrichment map plugin in Cytoscape (see material and methods). This network contains some well-known functions of Aurora-A including low- or moderately-enriched terms such as the mitotic cell cycle (4, 5, 9), apoptotic signalling pathway (48, 49), and viral life cycle (50) (**Figure 2A**). The high confidence (confidence = 0.9 out of 1.0) string-based protein-protein interaction network of the Aurora-A interactome (see materials and methods) also displays a dispersed cluster of proteins involved in the canonical function of Aurora-A in regulating mitotic cell cycle (**Supplementary Figure S2B**). This biological process includes well-characterized activators of Aurora-A such as TPX2 and CEP192 (51).

**Figure 2.**
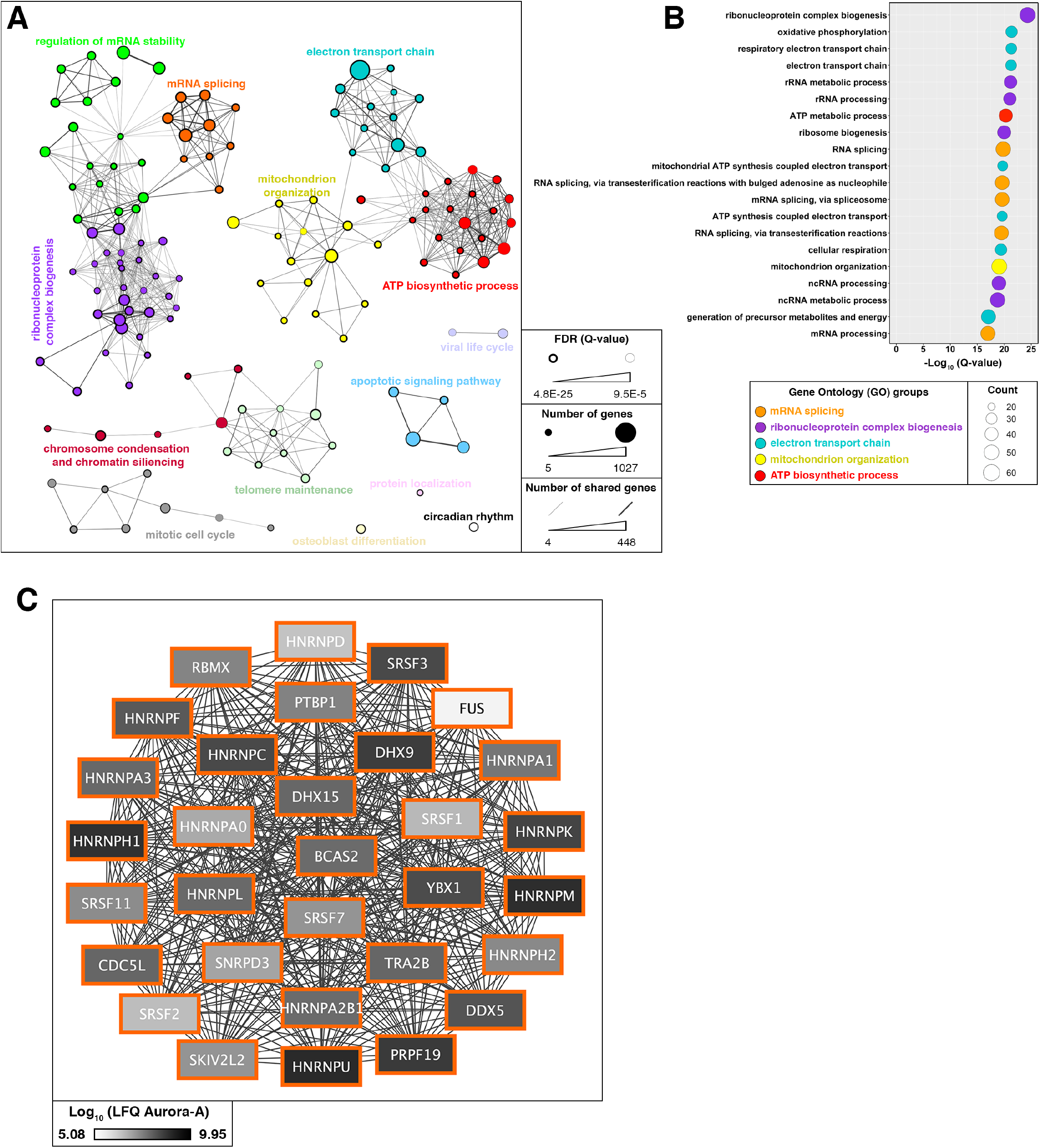
Gene enrichment analysis of the interactome revealing the new function of Aurora-A. (**A)** Enrichment map generated with Cytoscape after subjecting the Aurora-A interactome dataset to g:Profiler and annotating the Gene Ontology (GO) terms according to their function. (**B**) Dot plot showing the top 20 enriched GO term in g:Profiler indicating their significance and the number of genes (protein) counts that are mapped to each GO terms. (**C**) Cluster showing the string-based proteinprotein interaction of Aurora-A interacting proteins involved in mRNA splicing. The string-based proteinprotein interaction of all Aurora-A interacting proteins (407) and clusters (sub-networks) involved in other functions are shown in Supplementary Figure S2B.

The enrichment analysis further revealed few highly-enriched biological processes related to mitochondrial functions (**Figure 2A-B and Supplementary Figure S2A**), a novel function of Aurora-A that was described recently (11, 52, 53). Accordingly, string-based protein-protein interaction network of the Aurora-A interactome clearly contains clusters of proteins (sub-networks) that correspond to mitochondrial functions (**Supplementary Figure S2B**).

Notably predominant of the enriched terms are linked to RNA-related functions such as mRNA splicing, ribonucleoprotein complex biogenesis (ribosome biogenesis) and regulation of mRNA stability (**Figure 2A-B and Supplementary Figure S2A**). It has to be noted here that isolation of GFP-Aurora-A from protein cell extracts was performed in presence of nuclease, thereby minimising the chances of indirect co-immunoprecipitation of proteins via mRNA. The string-based protein-protein interaction network clearly reveals subnetwork corresponding to RNA splicing (**Figure 2C and Supplementary Figure S2B**). The gene enrichment analysis using two independent tools (g:Profiler and DAVID) also reflects RNA splicing as a significantly enriched biological process with at least 34 proteins corresponding to RNA splicing function (**Figure 2B and Supplementary Figure S2A**). Although it is so far unknown whether and how Aurora-A contributes to the regulation of RNA processing, two previous studies reported that Aurora-A affects RNA splicing by altering the stability of the splicing factor SRSF1 (ASF/SF2) (54, 55). Consistent with this observation, the repertoire of Aurora-A-associated proteins that we established contains several splicing factors, including SRSF1. These converging lines of experimental evidence led us to consider a potential connection between Aurora-A activity and RNA splicing.

### Aurora-A localizes to nuclear speckles

Pre-mRNA splicing proteins are enriched in specific intranuclear sites located in the interchromatin region of the nucleoplasm known as “nuclear speckles”. Under fluorescence microscope, nuclear speckles appear as irregularly shaped structures of varying size and are around 20 to 50 per cell depending on condition and cell type (56). As a first step toward unravelling a potential function of Aurora-A in RNA splicing, we investigated whether Aurora-A localizes in nuclear speckles through immunofluorescence microscopy experiments. U2OS cells transfected with GFP or GFP-Aurora-A expressing plasmids were stained with SC35 (SRSF2) antibodies, a nuclear speckle marker and GFP nanobodies to detect GFP (**Figure 3A**). Cells transfected with GFP-Aurora-A plasmid display punctuate patterns of GFP-Aurora-A in the nucleus and a marked co-localisation with SC35-stained nuclear speckles. By contrast, in cells transfected with GFP plasmid, a diffused pattern of GFP could be visualised with no particular GFP signal enrichment at SC35-stained nuclear speckles (**Figure 3A**).

**Figure 3.**
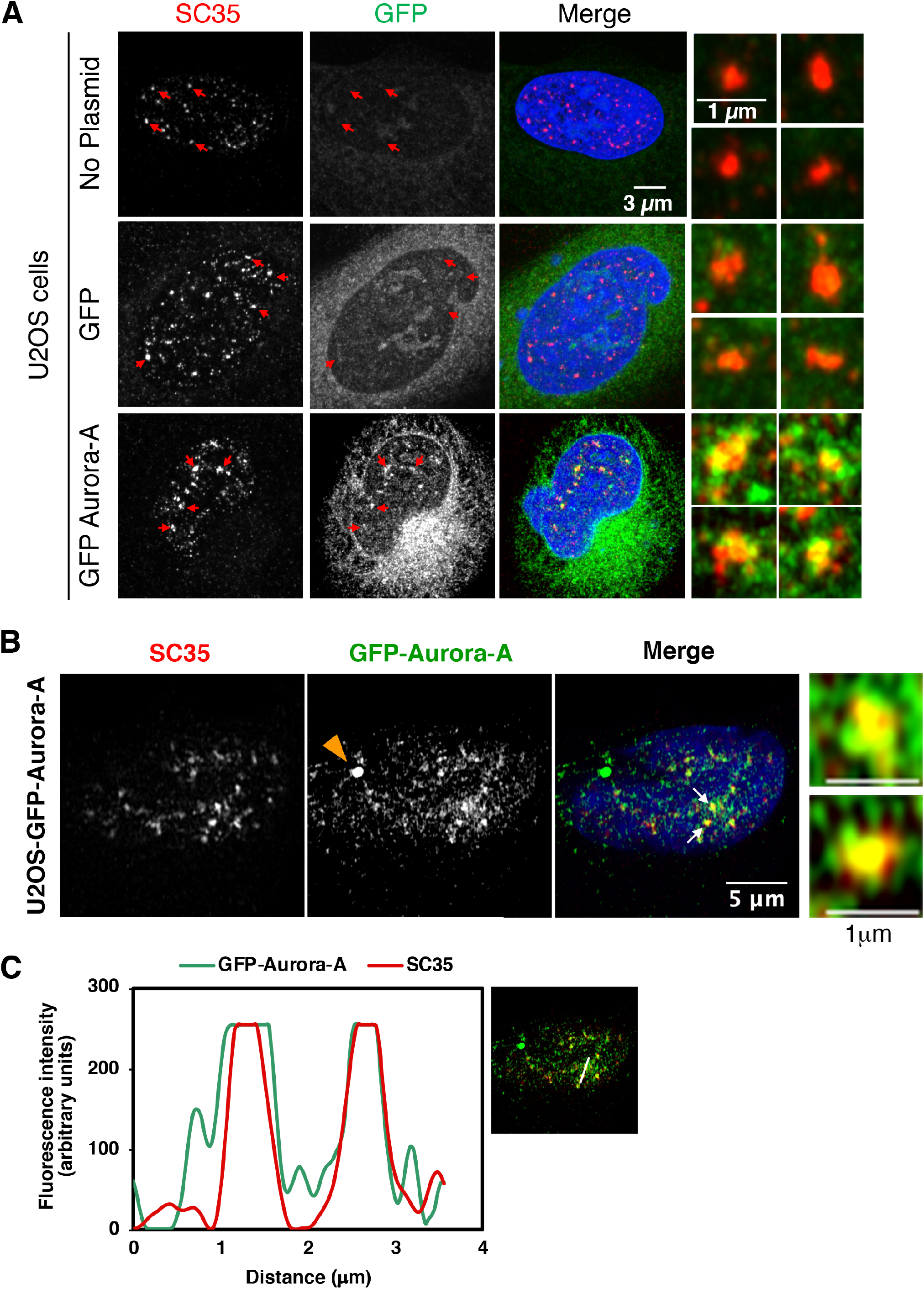
Aurora-A localizes to nuclear speckles. (**A)** Immunofluorescence microscopy of U2OS cells transfected with GFP and GFP-Aurora-A plasmids. Nuclear speckles were visualized with the anti-SC35 antibody (red) and DNA with DAPI (blue). Scale bar, 3 μm. The extreme right panels show enlarged versions of the regions indicated with red arrow in the first two left panels. Scale bar, 1 μm. (**B**) Immunofluorescence microscopy of U2OS cells stably expressing GFP-Aurora-A (green). Nuclear speckles were visualized with the anti-SC35 antibody (red) and DNA with DAPI (blue). The centrosome is indicated with an orange arrowhead in the left panel and two white arrows in the merged channel indicate the co-localization of Aurora-A with SC35. Scale bar, 5 μm. The extreme right panels show the magnification of the regions pointed with white arrow marks in the merged panels. Scale bar, 1 μm. The image represented here is from a single stack. (**C**) Line scan analysis using Fiji showing local intensity distributions of GFP-Aurora-A in green and SC35 in red. The merged image on the right indicates the position of the line scan with a white line.

To confirm this localization of Aurora-A at nuclear speckles and in order to exclude artefactual effect of plasmid over-expression, we performed similar experiment using U2OS cells that stably express GFP-tagged Aurora-A under the control of its own promoter. In this cell line as well, GFP-Aurora-A staining was again detectable in the form of punctuated pattern and showed co-localisation with SC35 staining (**Figure 3B**). Further, line scan analysis of nuclear speckles revealed an almost identical pattern between Aurora-A-GFP and SC35 signals (**Figure 3C**). Altogether, these observations demonstrate that Aurora-A kinase localizes to nuclear speckles and support the notion that Aurora-A participates in RNA splicing.

### Splicing proteins present in Aurora-A interactome predominantly correspond to non-core proteins involved in early steps of spliceosome assembly

The spliceosome comprises more than 200 proteins (21–23). To further characterize the interaction of Aurora-A with the spliceosome components, we took advantage of available proteomic data on the composition of spliceosome subcomplexes (22, 24, 25, 57). It has to be noted that the isolation of Aurora-A and its binding partners was performed in presence of nuclease and, therefore, in the absence of RNAs thus decreasing the chances of indirect interactions mediated by RNA molecules.

First, and consistent with our enrichment analysis (**Figure 2B and Supplementary Figure S2A**), the Aurora-A interactome presented a marked overlap with known splicing proteins, with more than one-sixth of previously reported spliceosomal proteins (22, 24, 25, 57) are part of Aurora-A interactome (**Figure 4A**). Notably, Aurora-A interacting splicing proteins belonged to distinct spliceosomal subcomplexes, including components of early assembly pre-E and E (commitment) complexes (hnRNP, SR proteins and mRNP), the presence of multiple subunits of same complexes reinforcing the reality of observed interactions. In addition, the presence of components of mid-assembly A and B complexes including components of U2 snRNP and Prp19 complex, and also potential components of catalytic C complexes as well as a few miscellaneous proteins and the common component of U snRNP (Sm proteins) could also be highlighted by this analysis (**Figure 4B**). Of note, subsets of proteins involved in early stages of spliceosome assembly (i.e. pre-E complex, E complex and A complex) are overrepresented (**Figure 4B**). Indeed, only 19.5% of the splicing proteins found in the Aurora-A interactome correspond to core component of the spliceosome involved in recognition of splicing sites and subsequent conformational changes leading to splicing catalysis (**Figure 4B**), including few DEAD/DEAH box helicases such as DDX5 and DHX15. By contrast, a vast majority (73.9%) of Aurora-A co-purified splicing proteins belonged to non-core components of the spliceosome, that is assumed to link the spliceosome to other cellular machineries (25). Remarkably, a substantial proportion (~56%) of these non-core proteins belong to two classes of trans-acting splicing factors called hnRNP and SR proteins that play regulatory roles on spliceosome, either promoting or preventing the recruitment of spliceosome at given RNA sites (58).

**Figure 4.**
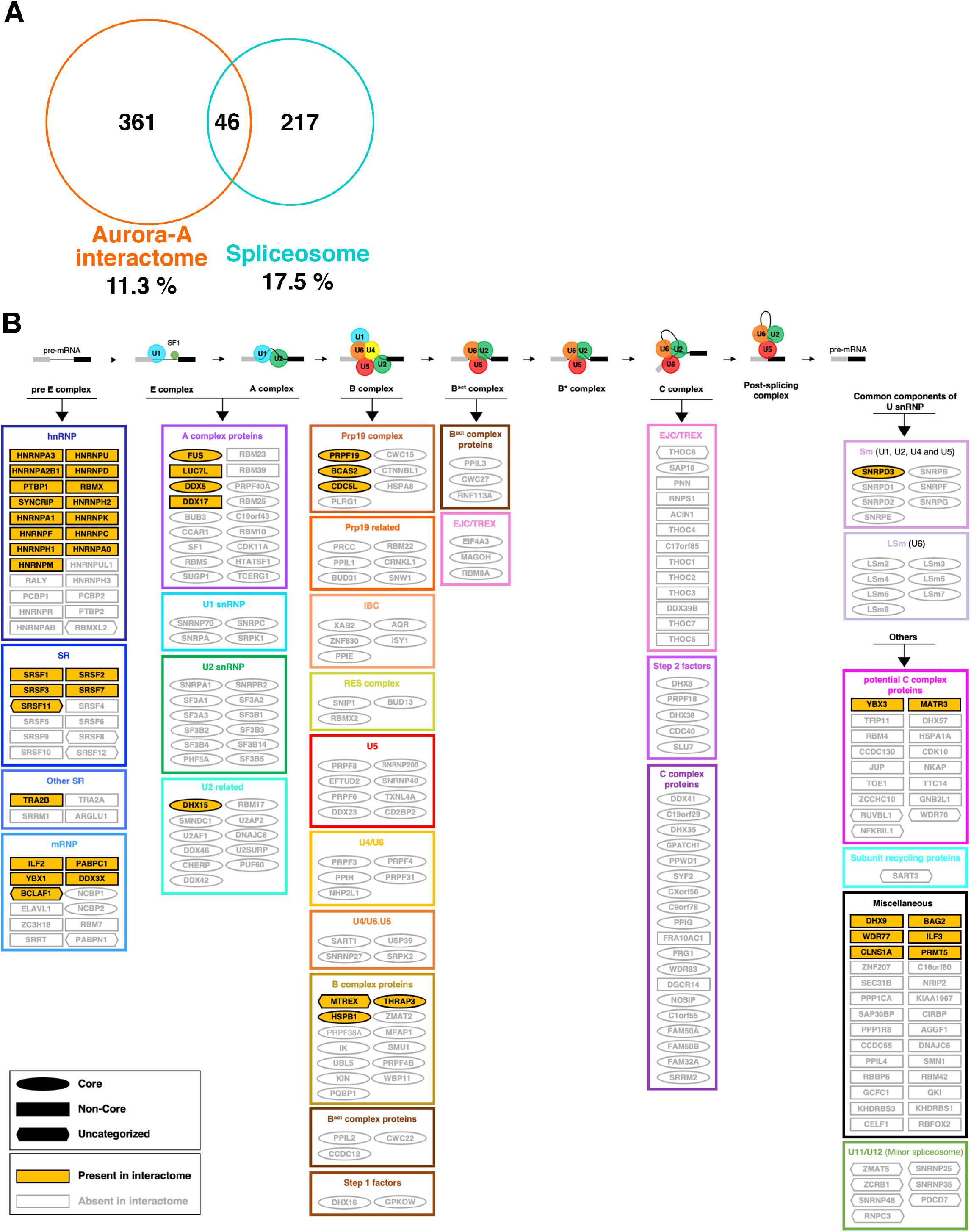
Interaction of Aurora-A with different components of the spliceosome. (**A)** Venn diagram showing splicing proteins interacting with Aurora-A by comparing Aurora-A interactome with the 263 known splicing proteins (22, 24, 25, 57). Numbers in the diagram refer to the number of proteins in each category. (**B**) Schematic drawing showing the categorization of Aurora-A interacting splicing partners into core and non-core component of the spliceosome. Each protein was categorized according to the spliceosomal sub-complexes in the order of their recruitment to the spliceosome.

Altogether, these results reveal a skewing of Aurora-A interactome towards non-core, regulatory splicing factors, suggesting that Aurora-A may impact on pre-mRNA splicing by modulating these splicing factors at early steps of splicing reaction.

### Aurora-A directly interacts with splicing factors, hnRNP and SR proteins *in vitro*

To determine whether proteins that are found in Aurora-A interactome interact with Aurora-A in a direct fashion, we employed *in vitro* pull-down assays. We first assessed the direct interaction of Aurora-A with three out of five canonical SR proteins found in the interactome dataset: SRSF1, SRSF3 and SRSF7. In that aim, purified recombinant GST-tagged SR proteins and HIS-tagged Aurora-A expressed in bacteria were incubated together, pulled down using glutathione affinity matrix and proteins recovered were subjected to western blot analysis using antibodies raised against Aurora-A. As shown in **Figure 5A**, Aurora-A was co-purified together with each of the three GST-tagged SR proteins but not with GST alone, indicating that Aurora-A is able to interact directly with SRSF1, SRSF3 and SRSF7 *in vitro*.

**Figure 5.**
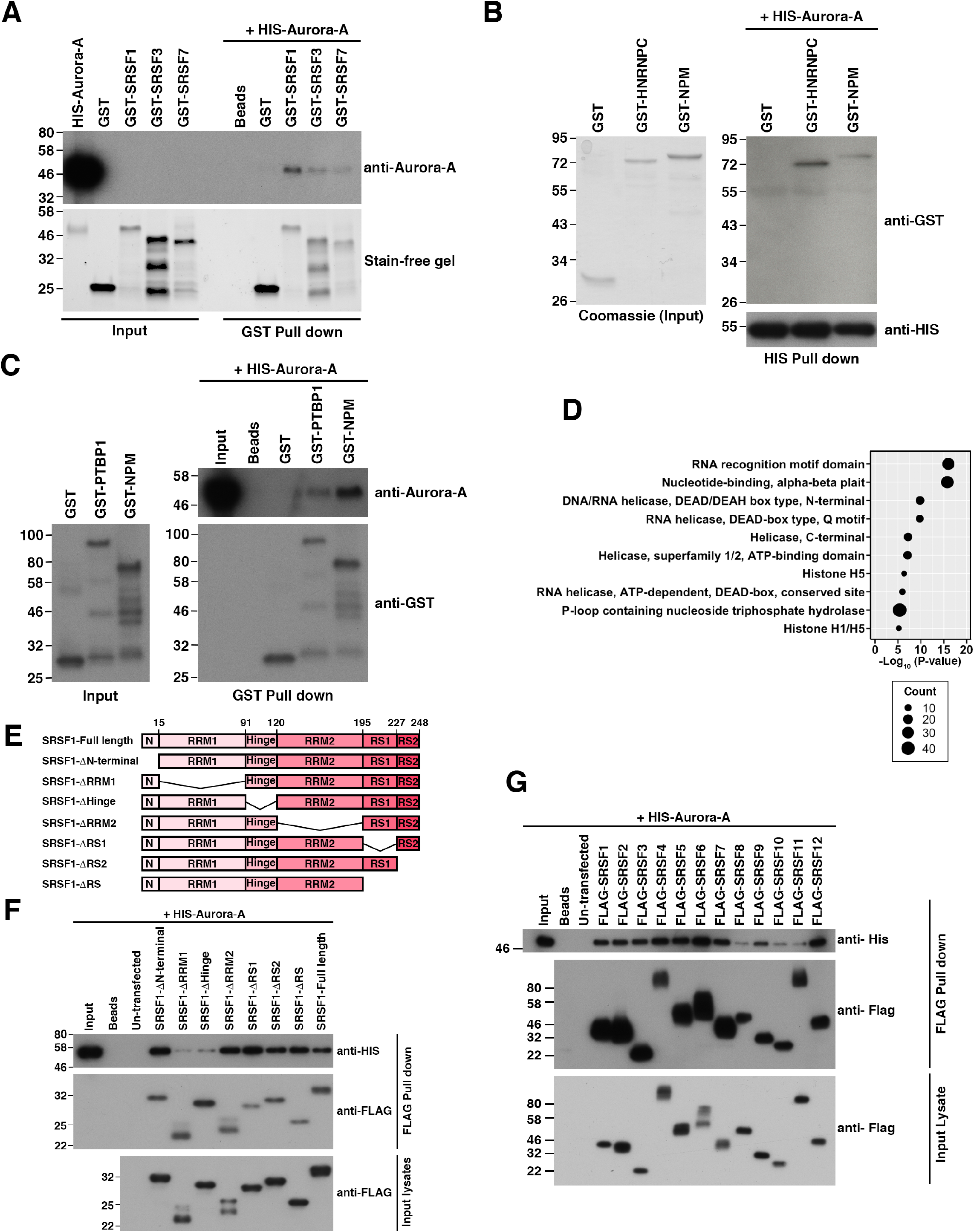
Aurora-A directly interacts with splicing factors. (**A**) GST pull-down assay showing the direct interaction of the SR proteins with Aurora-A *in vitro*. The bottom panel shows the stain-free gel image of purified recombinant HIS-tagged Aurora-A and GST-tagged SR proteins used for interaction assay. The top panel is the western blot showing the GST pulldown assay between HIS-tagged Aurora-A and GST-tagged SR proteins. GST was used as a negative control. (**B**) The HIS pull-down assay showing the direct interaction between HNRNPC protein and Aurora-A *in vitro*. The left panel shows the Coomassie staining of purified recombinant GST, GST-tagged HNRNPC and NPM used for interaction assay. The right panel is the western blot showing the HIS pull-down assay between HIS-tagged Aurora-A and GST-tagged proteins. GST was used as negative control and NPM was used as a positive control. (**C**) GST pull-down assay showing the direct interaction of PTBP1 with Aurora-A *in vitro*. The left panel shows the western blot of purified recombinant GST, GST-tagged PTBP1 and NPM used for interaction assay. The right panel is a western blot showing the GST pull-down assay between HIS-tagged Aurora-A and GST-tagged proteins. GST was used as negative control and NPM was used as a positive control. (**D**) Dot plot summarizing the domain enrichment analysis of Gene Ontology (GO) against the InterPro database using the filtered Aurora-A protein interaction dataset (407 Aurora-A interacting proteins). Top 10 predominant domain terms were plotted relative to their −log_10_(P values). (**E**) Schematic of full length and deletion mutants of FLAG-tagged SRSF1 protein used for FLAG pull-down assay in F. (**F**) FLAG pull-down assay showing the interaction of SRSF1 proteins with recombinant HIS-tagged Aurora-A *in vitro*. The bottom panel shows the western blot analysis of lysates from HEK293 cells that overexpress full-length and deletion mutants of FLAG-tagged SRSF1 proteins. The top two panels are western blot showing the interaction of Aurora-A with full-length and deletion mutants of SRSF1 proteins. (**G**) FLAG pull-down assay showing the direct interaction of the SR proteins with recombinant HIS-tagged Aurora-A *in vitro*. The bottom panel shows the western blot analysis of lysates from HEK293 cells that overexpress FLAG-tagged SRSF proteins. The top two panels are western blot showing the interaction of Aurora-A with SR proteins.

Next, we assessed the direct protein-protein interaction of Aurora-A with HNRNPC and PTBP1, two of the hnRNP proteins found in the interactome dataset. Purified recombinant HIS-tagged Aurora-A was incubated together with either GST-tagged HNRNPC (**Figure 5B**) or with GST-tagged PTBP1 (**Figure 5C**). Protein complexes were pulled down using cobalt column or glutathione matrix, respectively, and subjected to western analysis. Aurora-A pulled down GST-tagged HNRNPC and the positive control, NPM but not GST indicating that Aurora-A directly interacts with HNRNPC *in vitro* (**Figure 5B**) and, similarly, GST-tagged PTBP1 also co-pulled down Aurora-A (**Figure 5C**). Thus, a subset of splicing factors belonging to the hnRNP and SR protein families are direct interactors of Aurora-A *in vitro*.

Since it is well known that splicing factors share some well-characterised protein domains that mediate protein-protein or protein-RNA interactions (59), we hypothesized that one or more of these domains could mediate their direct interaction with Aurora-A. Therefore, we searched for such domains that would be enriched in the Aurora-A interactome using the DAVID tool. This domain enrichment analysis revealed significant enrichment for domains present in splicing factors, in particular the ‘RNA recognition motif’ domain (RRM), which was listed as the top enriched domain (**Figure 5D**). Remarkably, the same RRM motif also ranked first when enrichment analysis was restricted to the 46 Aurora-A interacting splicing proteins (**Supplementary Figure S3A**). Altogether, these domain enrichment analyses highlighted RRM domain as the major potential domain mediating interaction between splicing proteins and Aurora-A.

To test the possibility that RRM is actually involved in the interaction of splicing proteins with Aurora-A, we assayed the ability of versions of SRSF1 deleted for its different domains (**Figure 5E**) to interact with Aurora-A *in vitro*. In that goal, HIS-tagged Aurora-A was incubated with the different versions of FLAG-tagged SRSF1 expressed in and isolated from human HEK293 cells (**Figure 5F**). After incubation, proteins complexes were immunoprecipitated using anti-FLAG antibodies coupled to agarose beads (FLAG-M2 beads), eluted with FLAG peptides and subjected to western blot analysis. As shown in **Figure 5F**, HIS-tagged Aurora-A is pulled down by the full-length SRSF1 protein as well as by versions of SRSF1 deleted for N-terminal, RRM2, RS1 and RS2 domains. By contrast, versions that do not contain RRM1 or Hinge domains display markedly decreased amount of coimmunoprecipitated HIS-tagged Aurora-A. This domain analysis indicates that both the RRM1 and Hinge domain are required for the interaction of SRSF1 with Aurora-A *in vitro* (**Figure 5F**).

All the 12 canonical SR proteins are known to contain one or two RRM domains as well as RS domains (long repeat of Arginine and Serine dipeptides) (60). Building on the critical role of RRM domain we established for interacting with Aurora-A, we next tested the ability of all 12 canonical SR proteins to interact with Aurora-A *in vitro*. We performed *in vitro* FLAG pull-down assay using the FLAG-tagged SR (SRSF1-12) proteins and recombinant HIS-tagged Aurora-A, which revealed direct interaction of Aurora-A with all the 12 canonical SR proteins (**Figure 5G**), consistent with the role of RRM domain in mediating interaction with Aurora-A. RRM domains are present in 10 non-splicing and 22 splicing proteins from Aurora-A interactome, including HNRNPC, PTBP1, SRSF1, SRSF3 and SRSF7 for which the direct interaction with Aurora-A was validated (**Figure 5**). In agreement with these observations, Aurora-A also displayed direct interaction with RRM containing protein not present in our interactome such as RALY (**Supplementary Figure S3B**), further strengthening the relevance of RRM domain for Aurora-A interaction.

Altogether these results indicate that a number of splicing factors are bona fide interactors of Aurora-A and suggest that Aurora-A could contribute to the regulation of the biological functions of RRM-containing splicing factors.

### Aurora-A phosphorylates splicing factors *in vitro*

Since phosphorylation plays a crucial role during RNA splicing (61–65), we tested whether splicing factors that interact with Aurora-A *in vitro* are also a substrate of Aurora-A using *in vitro* kinase assays. Therefore, purified recombinant GST-tagged splicing factors were incubated with Aurora-A kinase *in vitro* in the presence of radiolabelled ^32^P-γATP. After incubation, proteins were separated by SDS-PAGE and radioactive signals were detected by autoradiography. As presented in **Figure 6**, all three SRSF proteins tested, but not GST alone, showed radioactive signals only in the presence of Aurora-A kinase, but not in its absence (**Figure 6A**). Likewise, HNRNPC and PTBP1 but not GST alone (**Figure 6B**), as well as RALY, PCBP2 and eIF4A3, but neither RS26 nor GST (**Supplementary Figure S3C**) display a radioactive signal in the presence of Aurora-A, indicating that the Aurora-A kinase phosphorylates splicing factors *in vitro*. Altogether, these results suggest that Aurora-A represents a novel splicing kinase that is able to phosphorylate several splicing factors found in complex with Aurora-A, possibly in order to regulate pre-mRNA splicing.

**Figure 6.**
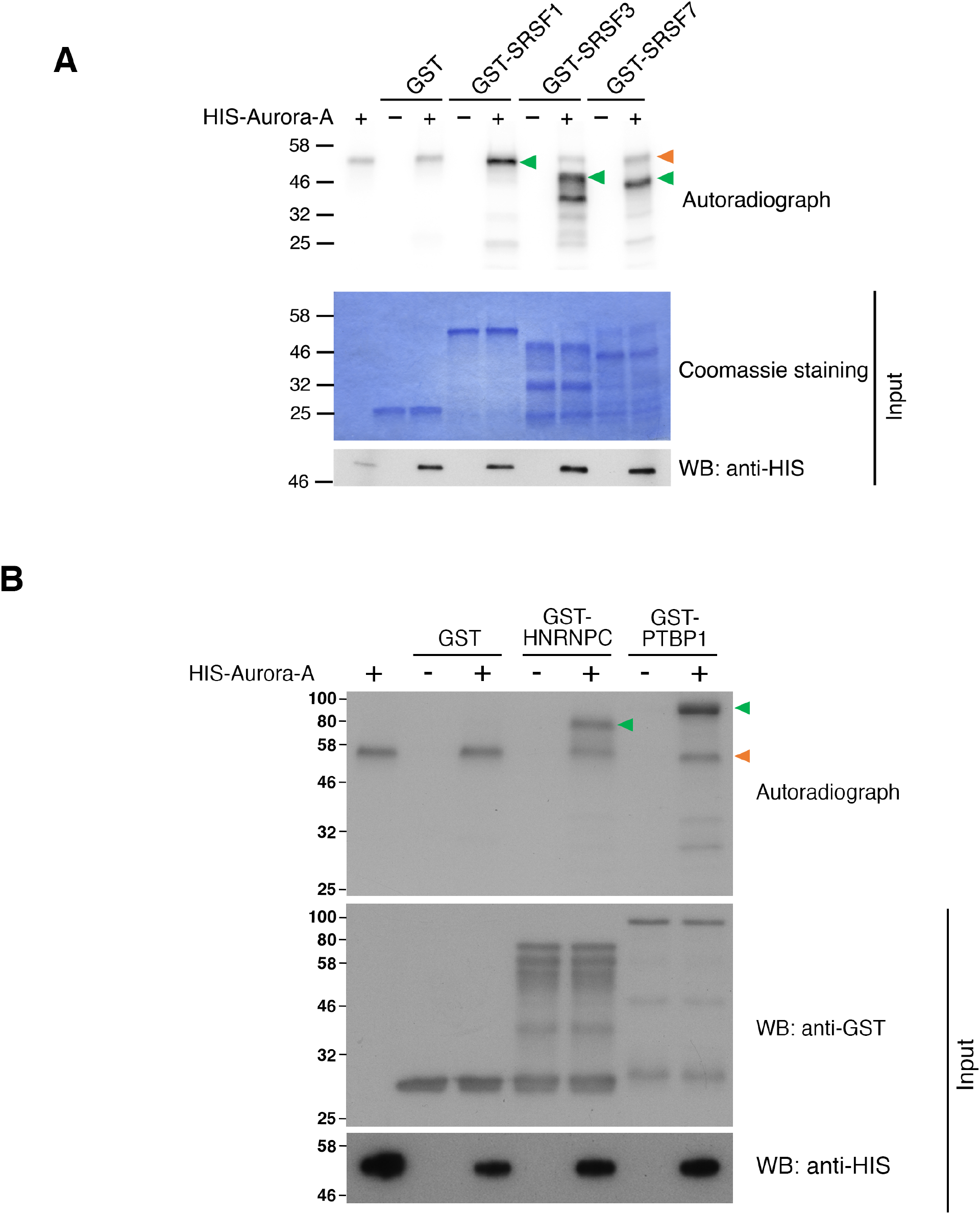
Aurora-A phosphorylates splicing factors *in vitro*. (**A)** Phosphorylation of SR proteins (SRSF1, SRSF3 and SRSF7) by Aurora-A was analysed by autoradiography on the top panel. GST was used as a negative control. Coomassie staining of purified recombinant GST-tagged SR proteins and western blot analysis of the HIS-tagged Aurora-A kinase are shown in the bottom two panels. Orange arrowhead indicates the auto-phosphorylation of Aurora-A and the green arrowhead indicates the phosphorylation of substrate (SR proteins). (**B**) Phosphorylation of hnRNP proteins (HNRNPC and PTBP1) by Aurora-A was analyzed by autoradiography on the top panel. GST was used as a negative control. Western blots of purified recombinant GST-tagged hnRNP proteins and HIS-tagged Aurora-A kinase are shown in the bottom two panels. Orange arrowhead indicates the auto-phosphorylation of Aurora-A and the green arrowhead indicates the phosphorylation of substrate (hnRNP proteins). GST was used as a negative control.

### Inhibition of Aurora-A modulates alternative splicing

Phosphorylation of splicing factors by specific kinases stands crucial for regulating their activities and biological functions (64). We have shown that Aurora-A interacts directly with trans-acting splicing factors and phosphorylates a subset of them *in vitro*, raising the interesting possibility that Aurora-A kinase activity is required for the modulation of alternative splicing. To test this hypothesis, we aimed at investigating whether and how inhibition of Aurora-A kinase activity affects pre-mRNA splicing in human cultured cells. However, the perturbation of Aurora-A is well known to affect cell cycle distribution by delaying mitotic entry, prolonging mitosis and inducing mitotic arrest (66–68). To circumvent this issue and, thereby, avoid the indirect effect of cell cycle perturbation on splicing changes, we coupled the inhibition of Aurora-A with cell synchronization procedure such that inhibitory treatment does not impact on cell cycle distribution (**Supplementary Figure S4A-C**). The results obtained from cells synchronised in G1, G2 and mitosis were pooled and analysed as a whole.

HeLa cells synchronisations were achieved by double thymidine arrest-release and were coupled to treatment with the Aurora-A pharmacological inhibitor MLN8237 for 24 hours, i.e. a complete cell cycle (see **Supplementary Figure S4A-C** for details). Synchronization of cell in mitosis was reached by further treating cells with spindle microtubule poison nocodazole. Proper synchronization of cells in G1, G2 and mitosis was assessed by FACS and by determination on mitotic indices, which revealed similar profiles between control and Aurora-A inhibited cells (**Supplementary Figure S4D-F**). Further, efficient inhibition of Aurora-A kinase activity in the different cell cycle series was assessed by immunoblotting experiments using an antibody that recognizes the phosphorylation of Aurora-A at threonine-288 residue, the activation mark of Aurora-A (**Supplementary Figure S4G-I**). 24 hours after Aurora-A inhibition, total RNAs were extracted and subjected to RNA sequencing analysis in order to analyse changes both in gene expression and in alternative splicing events that are caused by Aurora-A inhibition.

In total, VAST-TOOLS analysis (39, 40) identified 622 differentially regulated alternative splicing events (or 652 events including repetitive events across all cell cycle phases) upon Aurora-A inhibition with a threshold of ΔPSI > 15 or < −15% (**Supplementary Table S3**) while the edgeR tool (38) identified 60 differentially expressed gene (threshold of adjusted P-values of 0.05 and log_2_-fold change of 0.5, **Supplementary Table S4**). The 622 differentially spliced events occur in 505 genes. Notably, there was only one gene that is common between the differentially spliced genes and differentially expressed genes in Aurora-A inhibited condition, suggesting that alternative splicing and gene expression changes represent two separate layers of regulation upon inhibition of Aurora-A. It also underlines the fact that synchronization procedures allowed us to prevent indirect impact of Aurora-A inhibition on cell cycle progression. Furthermore, with a cut-off for log_2_-fold change of 1.5 and adjusted P-values of 0.05, only one gene (HIST2H2BE) was differentially expressed upon inhibition of Aurora-A (**Supplementary Table S4**). These results strongly indicate that under these experimental conditions Aurora-A inhibition has marginal effects on gene expression. Altogether our results suggest that the alternative splicing (AS) changes we identified are not a secondary effect of altered transcription but, instead, a direct consequence of Aurora-A inhibition on alternative splicing.

Out of the 622 AS changes affected by Aurora-A inhibition, 219, 199 and 234 AS event changes were detected at G1, G2 and mitotic phases respectively (**Figure 7A**), with only very little overlap between AS event changes across the different cell cycle populations (**Figure 7B**). With 49.7%, cassette exon was the major type of AS event changes detected followed by alternative 5’ splice site (Alt5’ SS) (20.4%) and alternative 3’ splice site (Alt3’ SS) (19.6%) whereas intron retention (IR) (10.3%) was a less common type of AS event changes (**Figure 7C and Supplementary Figure S5A-B**). When comparing the proportion of AS event changes due to Aurora-A inhibition with the total number of AS events detected, there appears to be no big bias towards any particular type of AS events (**Figure 7C**). Altogether, this result indicates that inhibition of Aurora-A activity globally impacts on the different types of AS events in a similar manner.

**Figure 7.**
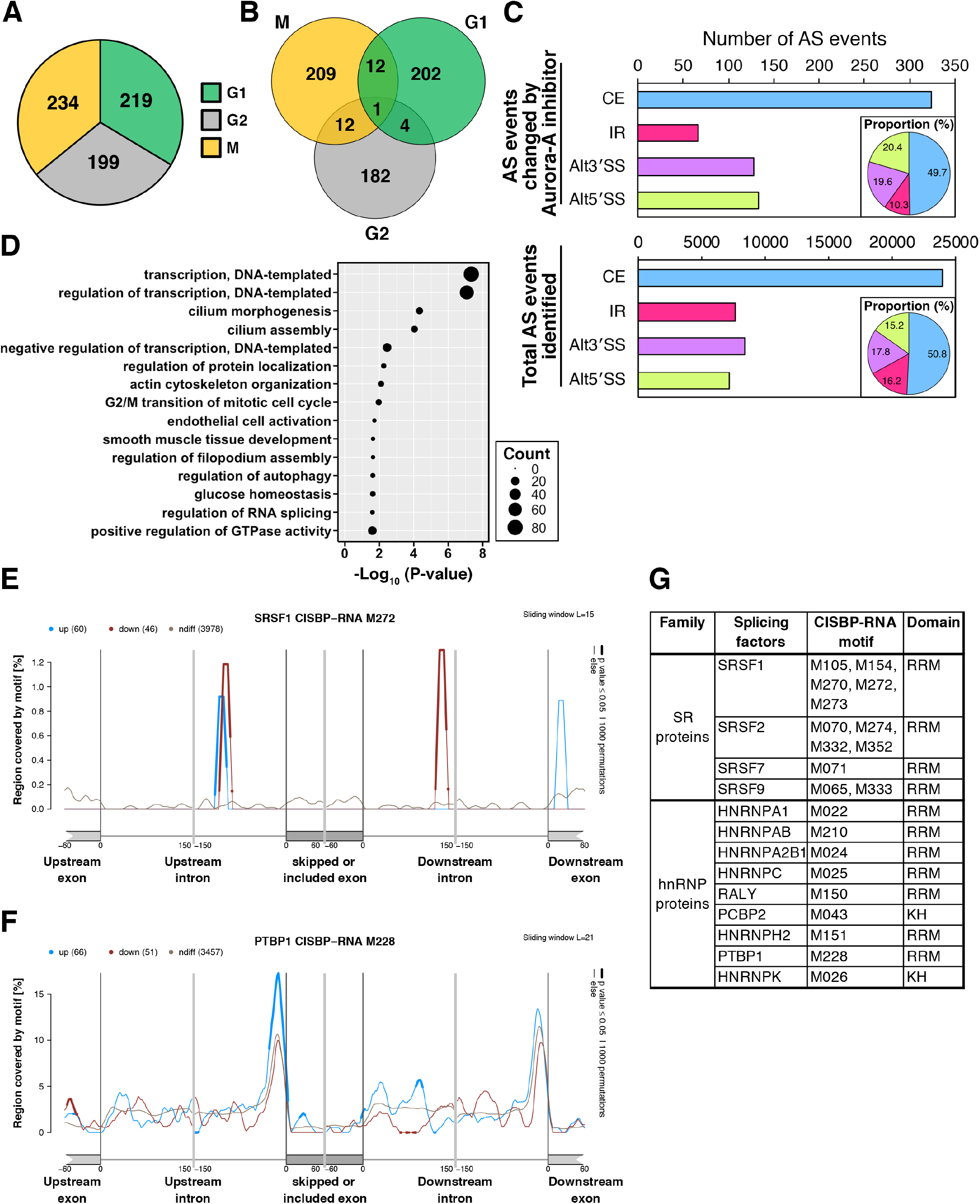
Aurora-A modulates the alternative splicing pattern. (**A)** Pie chart showing the distribution of differentially spliced events across cell cycle phases upon inhibition of Aurora-A. (**B**) Venn diagram showing the number of events differentially spliced at each cell cycle stage and the overlap of alternative splicing events across the cell cycle stages. (**C**) Bar graph showing the different categories of alternative splicing (AS) events affected by Aurora-A inhibition and the total number of AS events identified. The y-axis indicates the type of events and the x-axis indicates the number of differentially spliced events. The pie chart within the bar graph shows the distribution (proportion) of AS event types (**D**) Dot plot showing the top 15 enriched GO term obtained from the analysis of all the differentially spliced events across the cell cycle using the DAVID tool. The plot indicates the significance and the number of genes (protein) counts that are mapped to each GO terms. (**E**) Enrichment of RNA motif (M272) of SRSF1 in AS events affected by Aurora-A inhibition during mitosis. The blue, brown and grey lines show the enrichment for motif on upregulated (ΔPSI > 15%), downregulated (ΔPSI < −15%) and unregulated splicing events respectively. Regions with thick lines indicate the significant enrichment (FDR ≤0.05, 1000 permutation). (**F**) Enrichment of RNA motif (M228) of PTBP1 in AS events affected by Aurora-A inhibition in G2 phase. (**G**) Table summarising the RNA map results of the SR and hnRNP proteins.

Next, we aimed at determining whether mRNAs of which AS was affected by Aurora-A inhibition represent particular biological processes. Gene ontology analysis using DAVID tool for the 505 genes whose AS events altered upon Aurora-A inhibition revealed enrichment for biological processes such as regulation of transcription, cilium assembly, G2/M transition of the mitotic cell cycle, regulation of autophagy, glucose homeostasis, and positive regulation of GTPase activity (**Figure 7D**). Roles of Aurora-A in some of these functions have been reported previously (69–71), consistent with the possibility that Aurora-A regulates these functions at least in part by modulating splicing of concerned pre-mRNAs.

Finally, we examined the sequences of the pre-mRNAs, whose splicing is affected by the inhibition of Aurora-A, to search for RNA motifs to which splicing factors bind, in particular those we showed interacting directly with Aurora-A. To that aim, we used the Matt tool (41), which identified that particular RNA motifs for several SR and hnRNP proteins were indeed significantly enriched in the AS events altered by Aurora-A inhibition (**Figure 7E-F and Supplementary Figure S6-7**). For instance, the RNA motif M272, corresponding to sequences recognized by SRSF1, showed significant enrichment in the upstream intron of upregulated cassette exon events, whereas downregulated cassette exon events showed significant enrichment in the upstream and downstream introns (**Figure 7E)**. Similarly, for upregulated cassette exons, the RNA motif M228, corresponding to sequences recognized by PTBP1, showed significant enrichment in the 3’ region of the upstream intron, which includes the polypyrimidine tract (**Figure 7F)** that is known to mediate the regulatory functions of PTBP1 in some target genes (72). Notably, in agreement with the particular affinity of Aurora-A towards RRM domain-containing proteins, significant enrichment of RNA motifs of several RRM domain-containing proteins such as SRSF1, SRSF2, SRSF7, SRSF9, RALY, PTBP1, HNRNPA1, HNRNPH2, HNRNPA2B1 and HNRNPAB were observed (**Figure 7E-G and Supplementary Figure S6-7**). These results strongly suggest that regulation of the activity (e.g. binding) of these splicing factors contribute to mediate the AS changes identified upon pharmacological inhibition of the Aurora-A kinase.

Altogether, our work reveals that the kinase activity of Aurora-A regulates alternative splicing, likely through its interaction and phosphorylation of splicing factors hnRNP and SR protein families.

## DISCUSSION

### Interactome revealed novel functions of Aurora-A

Ever since the identification of Aurora-A as a potential drug target in cancer, many inhibitors have been developed, which unfortunately failed so far to clear the final stages of clinical trials due to their adverse effects (16). If Aurora-A is still to be considered as a promising anti-cancer target, it is essential to identify and understand its variety of cellular functions. The Aurora-A interactome we report here revealed novel interactors of Aurora-A, which suggested its involvement in ribosome biogenesis and RNA splicing. To our knowledge, Aurora-A has never been implicated in ribosome biology. However, two previous studies have associated Aurora-A with splicing, by showing that Aurora-A stabilizes SRSF1 and impacts on the splicing of apoptotic (55) and Androgen Receptor (54) genes. Remarkably, our Aurora-A interactome study identified 46 splicing proteins, suggesting a wider range of spliceosomal targets beyond SRSF1. This notion is further supported by the direct interaction and *in vitro* phosphorylation of multiple splicing factors (hnRNP and SR proteins) by Aurora-A. Given that hnRNP and SR proteins act as trans-acting splicing factors and affect splicing outcomes (73), we propose that Aurora-A modulates alternative splicing outcomes through interaction with and/or phosphorylation of these splicing factors.

### Functional implications of the interaction between Aurora-A and splicing proteins

We showed that Aurora-A interacts predominantly with non-core components of the spliceosome, in particular members of the hnRNP and SR protein families, which are regulatory factors of alternative splicing. Previous reports have identified the interaction of Aurora-A with hnRNP proteins such as hnRNP-K (12, 74) and hnRNP-U (75). Interaction of Aurora-A with hnRNP-K participates in MYC promoter trans-activation to enhance breast cancer stemness and in p53 activity regulation upon DNA damage (12, 74) while its interaction with hnRNP-U triggers its targeting to the mitotic spindle (75). However, the biological significance of these interactions in splicing has never been addressed. Our RNA-sequencing analysis had identified the AS event changes due to Aurora-A inhibition. Subsequently, RNA Maps analyses showed the enrichment for RNA motifs corresponding to sequences bound by Aurora-A-interacting SR and hnRNP proteins, thus strongly suggesting that regulation of these splicing factors by Aurora-A contributes to the identified AS changes. Therefore, our study has functionally linked the interaction between Aurora-A and splicing factors to RNA splicing.

RRM domain of splicing factors was initially thought to be important for their interaction to RNA but it was later identified to also facilitate protein-protein interactions (76, 77). We have shown evidence that RRM1 but not of RRM2 of SRSF1 is essential for its interaction with Aurora-A as RRM1 represents the canonical form of RRM whereas the RRM2 is an atypical or pseudo-RRM (78). Similar interaction to canonical RRM but not to pseudo-RRM has been observed for MDM2 and RPL5 (79). This observation strongly supports the notion that canonical RRM but not pseudo-RRM domain mediates direct physical interaction between splicing factors and Aurora-A.

To a lesser extent, Aurora-A also interacted with core spliceosomal proteins such as the RNA helicases (DDX5 and DHX15) and PRP19 complex (PRPF19, BCAS2 and CDC5L). The RNA helicases facilitate structural and conformational remodelling between the snRNAs at different stage of splicing reaction (80–83). Next to RRM, RNA helicase domain was the most enriched domain in our enrichment analysis. The PRP19 complex is important for the activation of spliceosome and formation of stable tri-snRNP (84, 85). The RNA helicases, PRP19 complex and splicing factors are also implicated in other gene regulatory mechanisms such as gene expression, RNA export, RNA degradation, Ribosome biogenesis and translation (86–89). Therefore, it remains possible that Aurora-A participates in these splicing and other gene regulatory mechanisms through the regulation of these splicing proteins.

### Aurora-A is a novel splicing kinase

Phosphorylation regulates multiple crucial steps during pre-mRNA splicing. It influences dynamic structural reorganization of the spliceosome by modulating protein-protein interactions, recruitment of splicing factors from the nuclear speckles to the site of active transcription, the catalysis of the splicing reaction and also the splicing outcomes (62–64, 90). In our work, we showed that Aurora-A phosphorylates several SR and hnRNP proteins *in vitro*. A previous phospho-proteomics study has identified the splicing factors, HNRNPA1, HNRNPK and RBMX, which were also present in our interactome, as potential *in vivo* substrates of Aurora-A (91). Aurora-A may modulate alternative splicing through phosphorylation of splicing factors. In agreement with this hypothesis, we observed that inhibition of its kinase activity perturbs alternative splicing. Thus, our work introduces Aurora-A as a new member in the family of splicing regulating kinases. Similar to Aurora-A, other kinases (AKT, MAPKs, PTKs, NEK2 and PKA) whose normal functions involve signal transduction and cell cycle regulation, also influence splicing mechanisms (92–97). These kinases could help the cells to modulate the splicing mechanism in response to physiological and pathological stimuli and thereby link the splicing mechanism with other cellular processes.

### Effect of Aurora-A inhibition on alternative splicing and their impact on cancer

Pharmacological inhibition of Aurora-A kinase activity resulted in altered patterns of 622 AS events. In contrast to previous studies (54, 55), we did not detect alteration in the splicing of either apoptotic genes or the AR gene. We, therefore, postulate that the effect of Aurora-A on the splicing of these genes could be cell context-dependent. For instance, the effect of Aurora-A on the splicing of the AR gene could be specific to prostate cancer cells. Indeed, we did detect a decrease in SRSF1 levels due to Aurora-A inhibition in mitotic cells (**Supplementary Figure S8**), in agreement with previous studies (54, 55). We believe that the experimental conditions such as the dose of Aurora-A inhibitor and cell synchronization procedures that were used in our study do not favour pro-apoptotic splicing. This is in agreement with the observation that the pro-apoptotic splicing changes are observed in a dosedependent manner (55).

Further, we have evidenced that Aurora-A inhibition affects alternative splicing of genes involved in cell cycle-related functions such as cilium assembly, GTPase activity and G2/M transition. These cell cycle-related functions are known to be mis-regulated in cancer (98–101). It is thus conceivable that Aurora-A assists cancer cell growth by regulating these cell cycle-related processes at least in part through modulation of alternative splicing beside its well-established cell cycle functions.

### Exploiting the interaction of Aurora-A with splicing proteins for developing cancer therapeutics

RNA splicing and cell cycle are strongly interconnected. For instance, depletion of several splicing proteins leads to hampering of cell cycle progression (102–105). Conversely, the SR protein, SRSF10 in its dephosphorylated form, represses splicing during mitosis (106). Additionally, perturbation of several cell cycle regulators such as NEK2 and Aurora-A impacts on RNA splicing (54, 55, 92). Often these splicing and cell cycle regulating kinases are mis-regulated in diseases such as cancer (64, 107–109). Since these non-canonical splicing kinases are also involved in other splicing independent normal functions, it remains an important challenge to selectively target the splicing function of these kinases in cancer. In our interactome study, the domain enrichment analysis revealed the particular affinity of Aurora-A towards specific domains, in particular the RRM domain present in numerous splicing factors. Therefore, in the case of Aurora-A, its splicing function could be specifically targeted by modulating the affinity of Aurora-A towards RRM domains. In the end, we believe that understanding of all the new functions of Aurora-A and targeting each of its tumorigenic functions individually might help to overcome toxic side effects when targeting Aurora-A in cancer.

## DATA AVAILABILITY

The mass spectrometry proteomics data of Aurora-A have been deposited to the ProteomeXchange Consortium (110) via the PRIDE (111) partner repository with the dataset identifier PXD021093.

RNA-sequencing data generated from this study have been deposited in Gene Expression Omnibus (112) under the accession number GSE160733.

## ACKNOWLEDGEMENTS

We are grateful to Stephanie Dutertre of the Microscopy-Rennes Imaging Center (Biologie, Santé, Innovation Technologique, BIOSIT, Rennes, France). We thank CRG Genomic Unit for sequencing our samples. We thank Dr. Thomas Gonatopoulos-Pournatzis (National Cancer Institute, USA) for the constructive comments. We thank Kanchana Nandanan Chathoth (CIMIAD, U1241 NUMECAN, Rennes, France) for proof reading the article.

