## Supplementary for "Aurora-A phosphorylates splicing factors and regulates alternative splicing"

† The authors wish to be known that, in their opinion, the last two authors should be regarded as Joint Last Authors.

#### SUPPLEMENTARY FIGURES AND TABLES LEGENDS

**Supplementary Figure S1. U2OS cells expressing GFP-Aurora-A and Aurora-A-shRNA displays normal localization and cell cycle.**

**(A)** Snapshots of lifetime imaging of GFP-Aurora-A localization in U2OS-GFP-Aurora-A-shRNA-Aurora-A stable cell during interphase and mitosis. GFP-Aurora-A is shown in green and DNA in blue. Scale bar: 10 $\mu$m. GFP-Aurora-A is localized to the centrosome and mitotic spindles at interphase and mitosis respectively. **(B)** Cell cycle distribution of wild-type U2OS compared to U2OS expressing GFP-Aurora-A and Aurora-A-shRNA. The cell cycle distribution was monitored by FACS analysis.

**Supplementary Figure S2. Protein-protein interaction network of Aurora-A interacting proteins**

**(A)** Dot plot showing the top 20 enriched GO term obtained from the interactome data using the DAVID tool. The plot indicates the significance and the number of genes (protein) counts that are mapped to each GO terms. (**B**) The string-based protein-protein interaction created using Cytoscape showing the clusters (sub-networks). The proteins involved in mRNA splicing, ribonucleoprotein complex biogenesis and electron transport chain form dense clusters whereas proteins involved in mitochondrial organization and mitotic cell cycle form dispersed clusters.

**Supplementary Figure S3. Aurora-A interacts with and phosphorylates splicing factors *in vitro***

(**A**) Dot plot summarizing the domain enrichment analysis of Gene Ontology (GO) against the InterPro database using the splicing factors interacting with Aurora-A (46 proteins). Top predominant domain terms were plotted relative to their -$\log_{10}$(P values). (**B**) GST pull-down assay showing the direct interaction of the splicing proteins with Aurora-A *in vitro*. The left panel shows the Coomassie staining of purified recombinant HIS-tagged Aurora-A and GST-tagged splicing proteins used for interaction assay. The right panel is the western blot showing the GST pull-down assay between HIS-tagged Aurora-A and GST-tagged splicing proteins. GST was used as negative control and NPM was used as a positive control. (**C**) Phosphorylation of splicing proteins by Aurora-A analyzed by autoradiography. The orange arrowhead indicates the auto-phosphorylation of Aurora-A and green arrowhead indicates the phosphorylation of substrates (splicing proteins). GST was used as negative control and GST-H3 was used as the positive control.

**Supplementary Figure S4. Inhibition of Aurora-A in synchronized HeLa cells.**

**(A)** The scheme shows the condition used for synchronizing HeLa cells at G1 phase using double thymidine block. Mitotic cells were removed by mitotic shake-off just before harvesting the cells. On red is the cell cycle phases through which Aurora-A remain inhibited (**B**) The scheme shows the condition used for synchronizing HeLa cells at G2 phase after releasing cells from double thymidine block for 6 hours. Mitotic cells were removed by mitotic shake-off just before harvesting the cells. On red is the cell cycle phases through which Aurora-A remain inhibited (**C**) The scheme shows the condition used for synchronizing HeLa cells at mitotic phase by releasing cells from double thymidine block for 12 hours. Nocodazole was used in the last two hours of synchronization to prevent the release of cells from mitosis. Mitotic cells were collected by mitotic shake-off. On red is the cell cycle phases through which

Aurora-A remain inhibited (**D**) The graph showing cell cycle profile from FACS indicating the synchronization of HeLa cells at G1 phase. (**E**) The graph showing cell cycle profile from FACS and mitotic index indicating the synchronization of HeLa cells at G2 phase. (**F**) The graph showing cell cycle profile from FACS and mitotic index indicating the synchronization of HeLa cells at mitotic phase. (**G**) Western blot analysis of HeLa cells synchronized at G1 phase showing the decrease in the levels of Thr228 phosphorylation of Aurora-A. (**H**) Western blot analysis of HeLa cells synchronized at G2 phase showing the decrease in the levels of Thr228 phosphorylation of Aurora-A. (**I**) Western blot analysis of HeLa cells synchronized at mitotic phase showing the decrease in the levels of Thr228 phosphorylation of Aurora-A.

**Supplementary Figure S5. Distribution of different types of spliced events upon inhibition of Aurora-A.**
(**A**) Bar graph showing the different categories of alternative splicing events that are either towards more inclusion ($\Delta$PSI < -15%) or more skipping ($\Delta$PSI > 15%) by Aurora-A inhibition. (**B**) Bar graph showing the different categories of alternative splicing (AS) events affected by Aurora-A inhibition in G1, G2 and mitotic phases. The y-axis indicates the type of events and the x-axis indicates the number of differentially splicing events. The pie chart within the bar graphs shows the distribution (proportion) of AS event types.

**Supplementary Figure S6. RNA map motif analysis of SR proteins.**
Enrichment of RNA motifs of SR proteins in AS events affected by Aurora-A inhibition. The blue, brown and grey lines show the enrichment for motif on upregulated ($\Delta$PSI > 15%), downregulated ($\Delta$PSI < -15%) and unregulated splicing events respectively. Regions with thick lines indicate the significant enrichment (FDR ≤0.05, 1000 permutation).

**Supplementary Figure S7. RNA map motif analysis of hnRNP proteins.**
Enrichment of RNA motifs of hnRNP proteins in AS events affected by Aurora-A inhibition. The blue, brown and grey lines show the enrichment for motif on upregulated ($\Delta$PSI > 15%), downregulated ($\Delta$PSI < -15%) and unregulated splicing events respectively. Regions with thick lines indicate the significant enrichment (FDR ≤0.05, 1000 permutation).

**Supplementary Figure S8. Effect of Aurora-A inhibition on ASF/SF2 (SRSF1) proteins levels.**
(**A**) Western blot analysis of HeLa cells synchronized at G1, G2 and mitotic phases showing the levels of ASF/SF2 (SRSF1) upon Aurora-A inhibition.

**Supplementary Table S1.** Excel sheet showing the list of primers used in this study.

**Supplementary Table S2.** Excel workbook showing the interacting partners of Aurora-A identified by affinity-purification coupled to mass spectrometry (AP-MS). The workbook also includes the list of common contaminants and unfiltered proteomics datasets.

**Supplementary Table S3.** Excel workbook showing differentially spliced events upon Aurora-A inhibition. The worksheets were labelled according to the cell cycle stages (G1, G2 and M).

**Supplementary Table S4.** Excel workbook showing differentially expressed genes upon Aurora-A inhibition.

### Supplementary Figure-S1

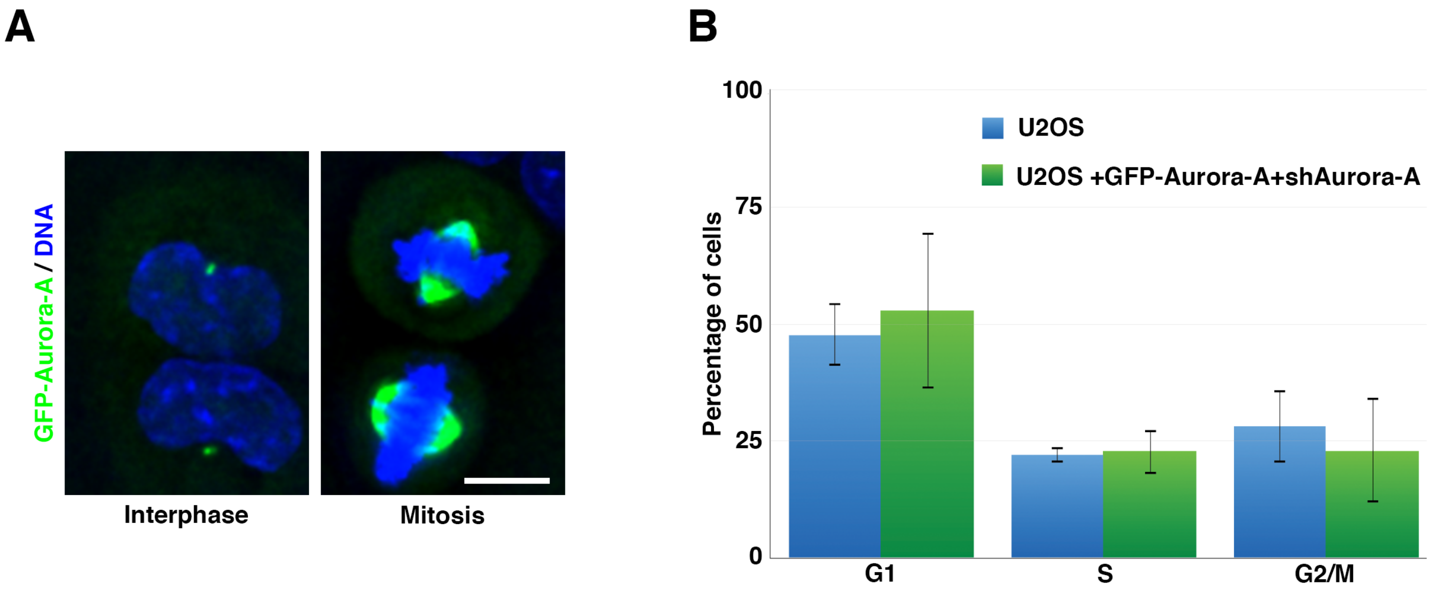

### Supplementary Figure-S2

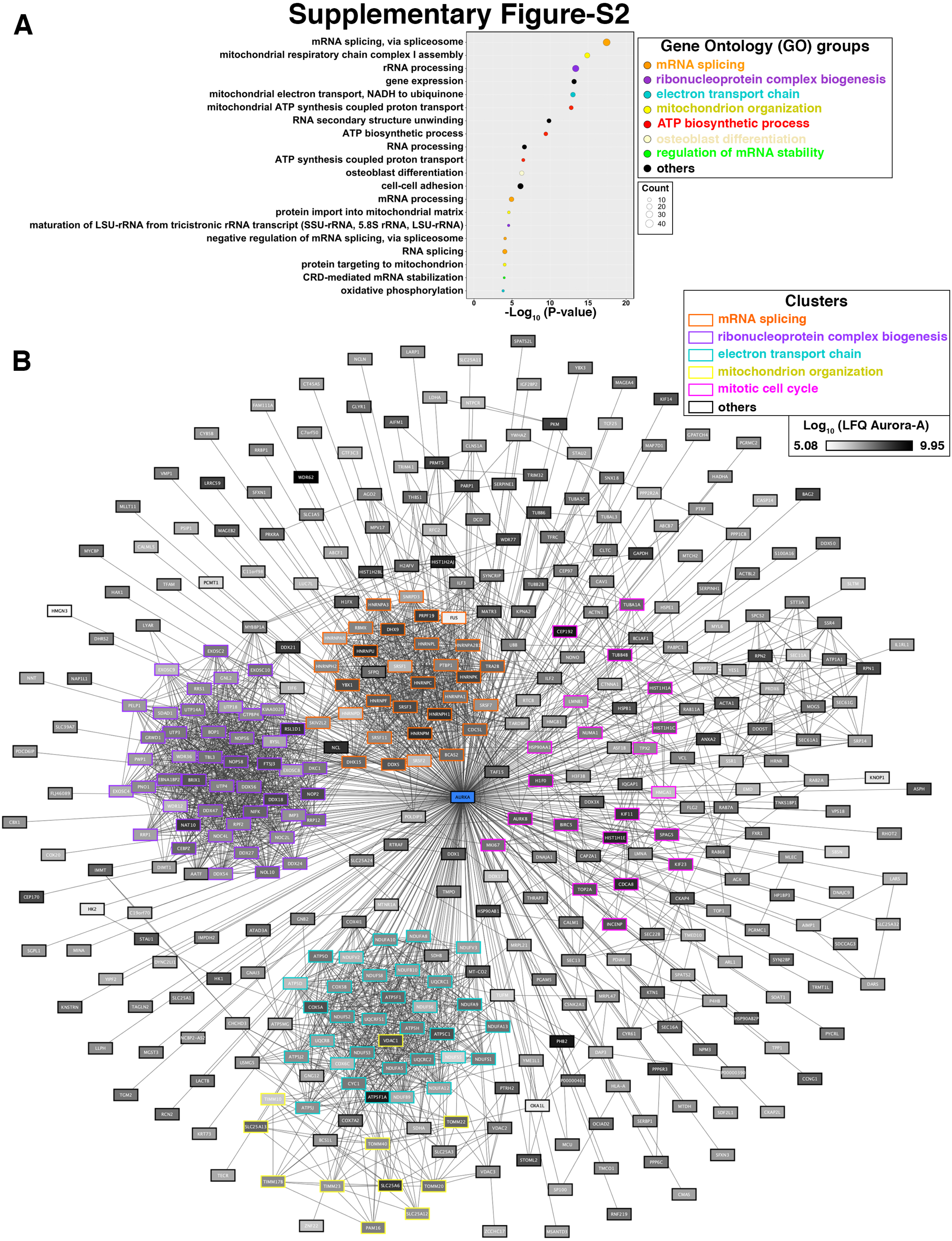

### Supplementary Figure-S3

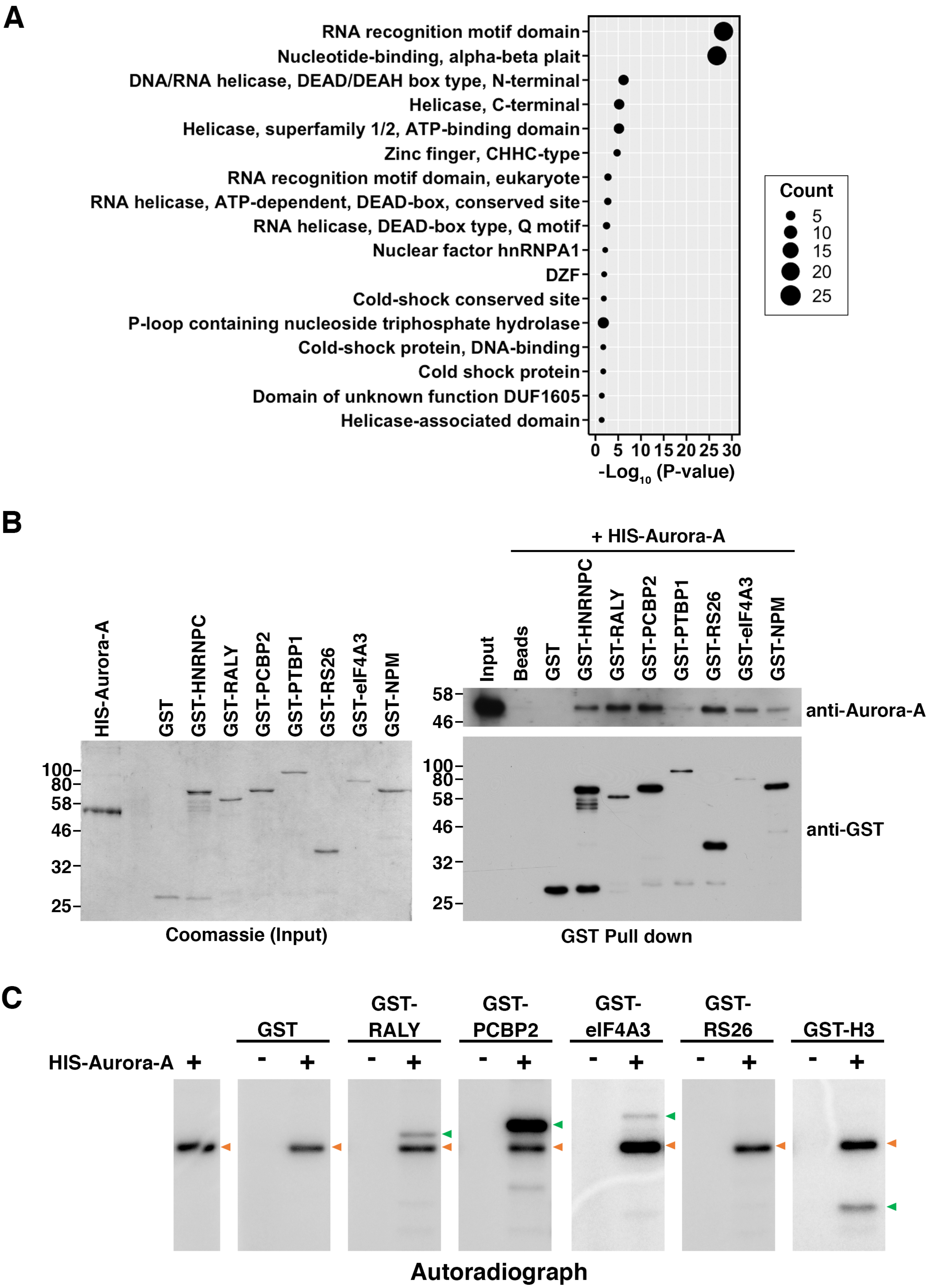

### Supplementary Figure-S4

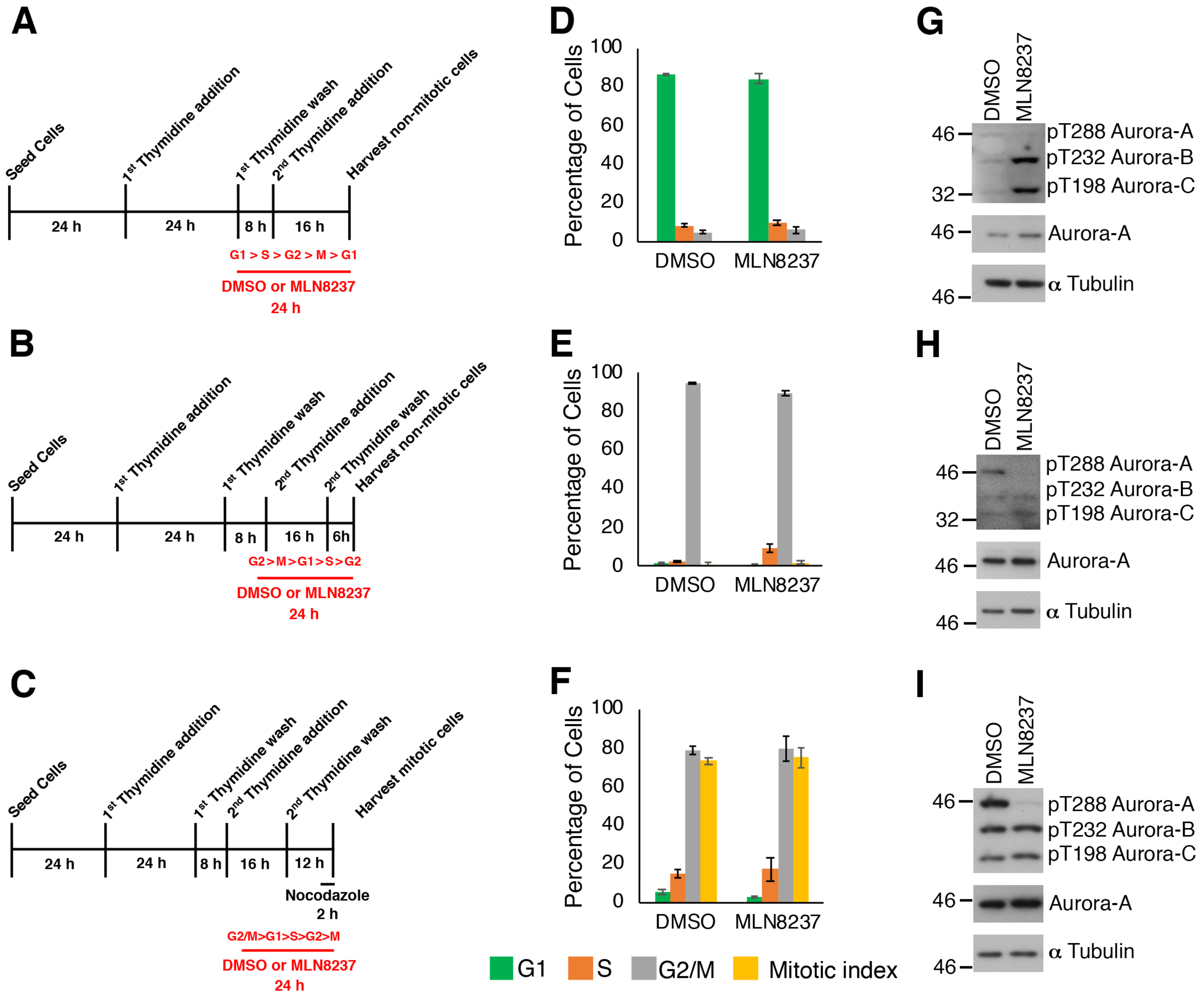

### Supplementary Figure-S5

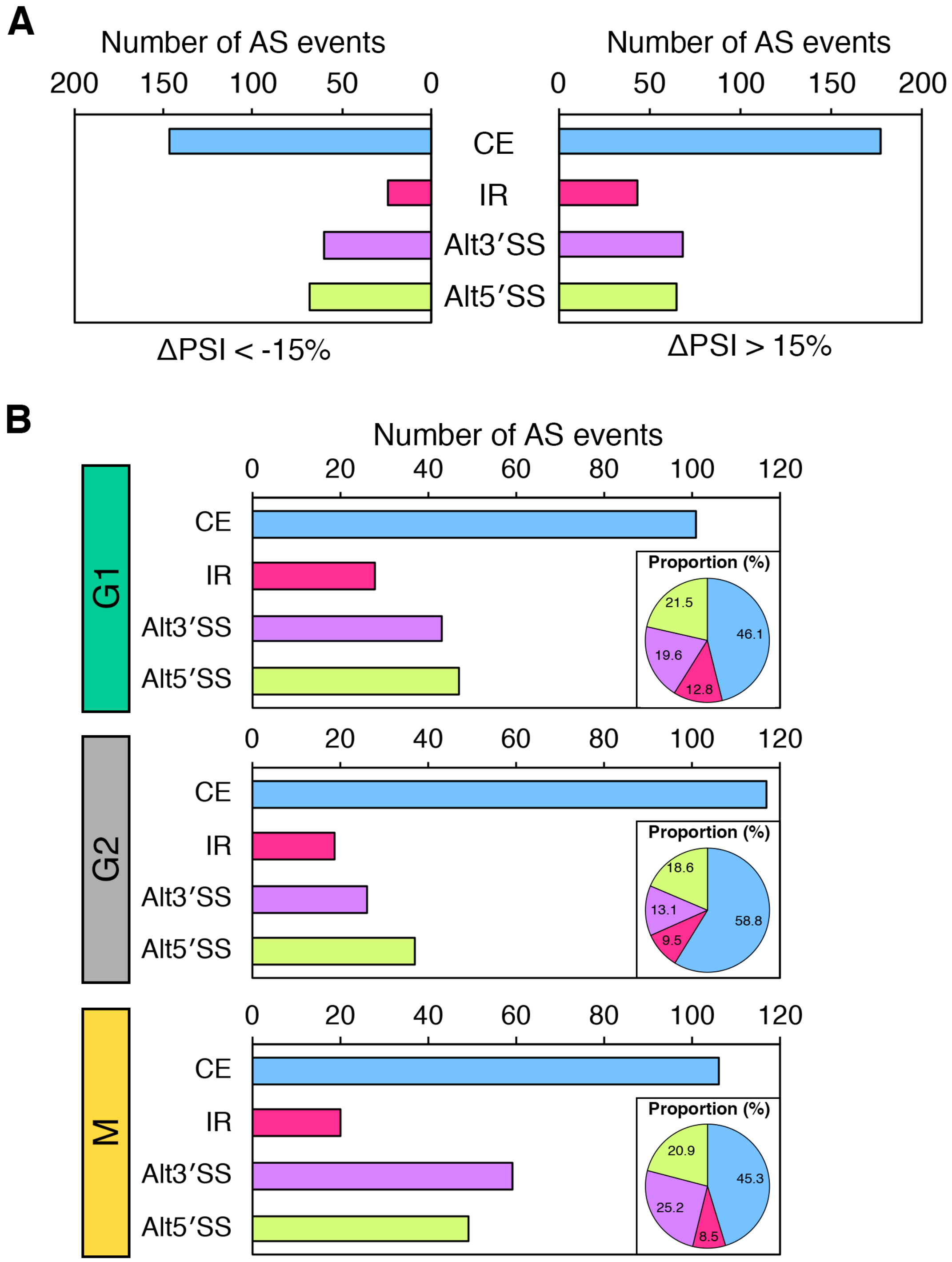

### Supplementary Figure-S6

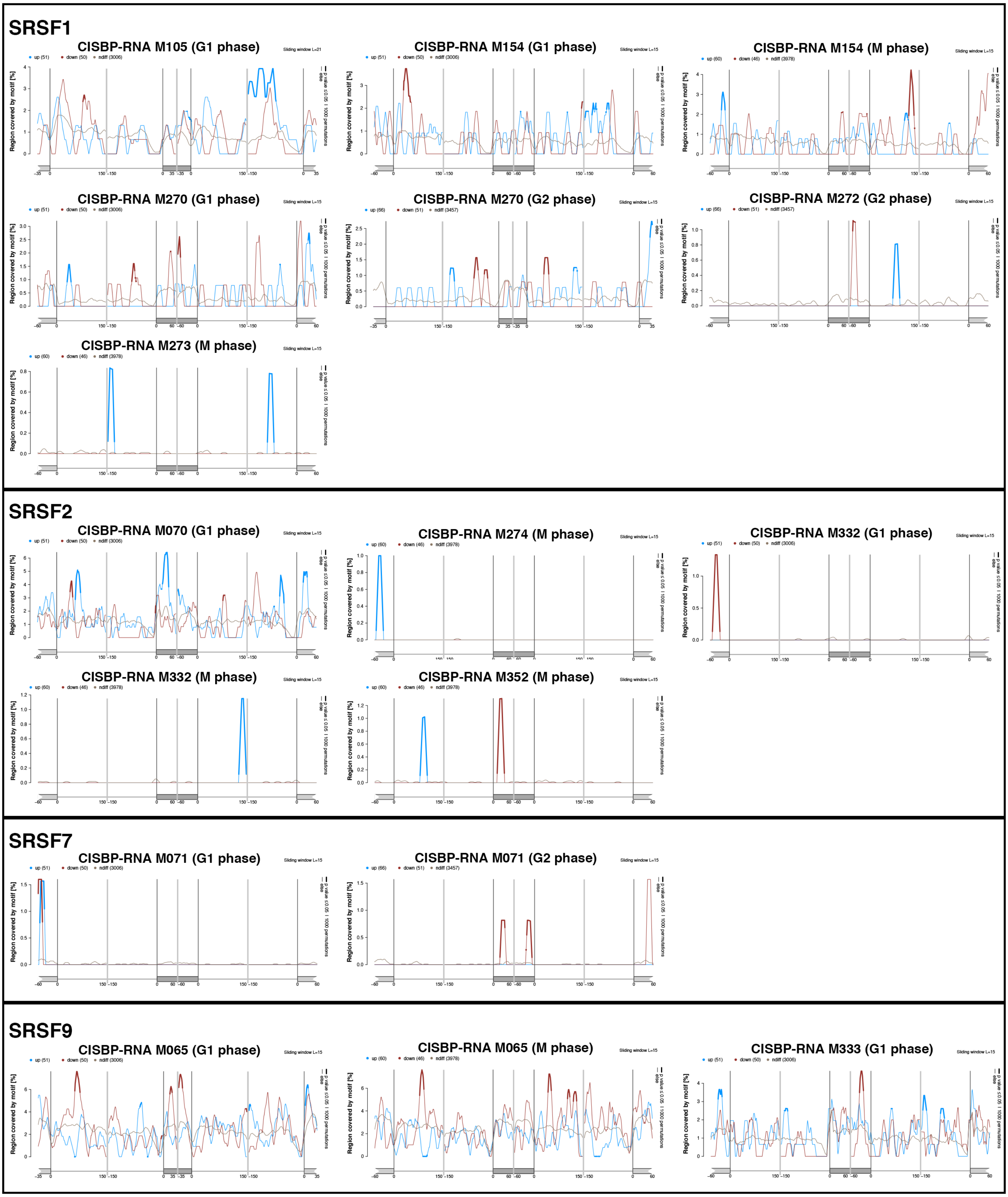

### Supplementary Figure-S7

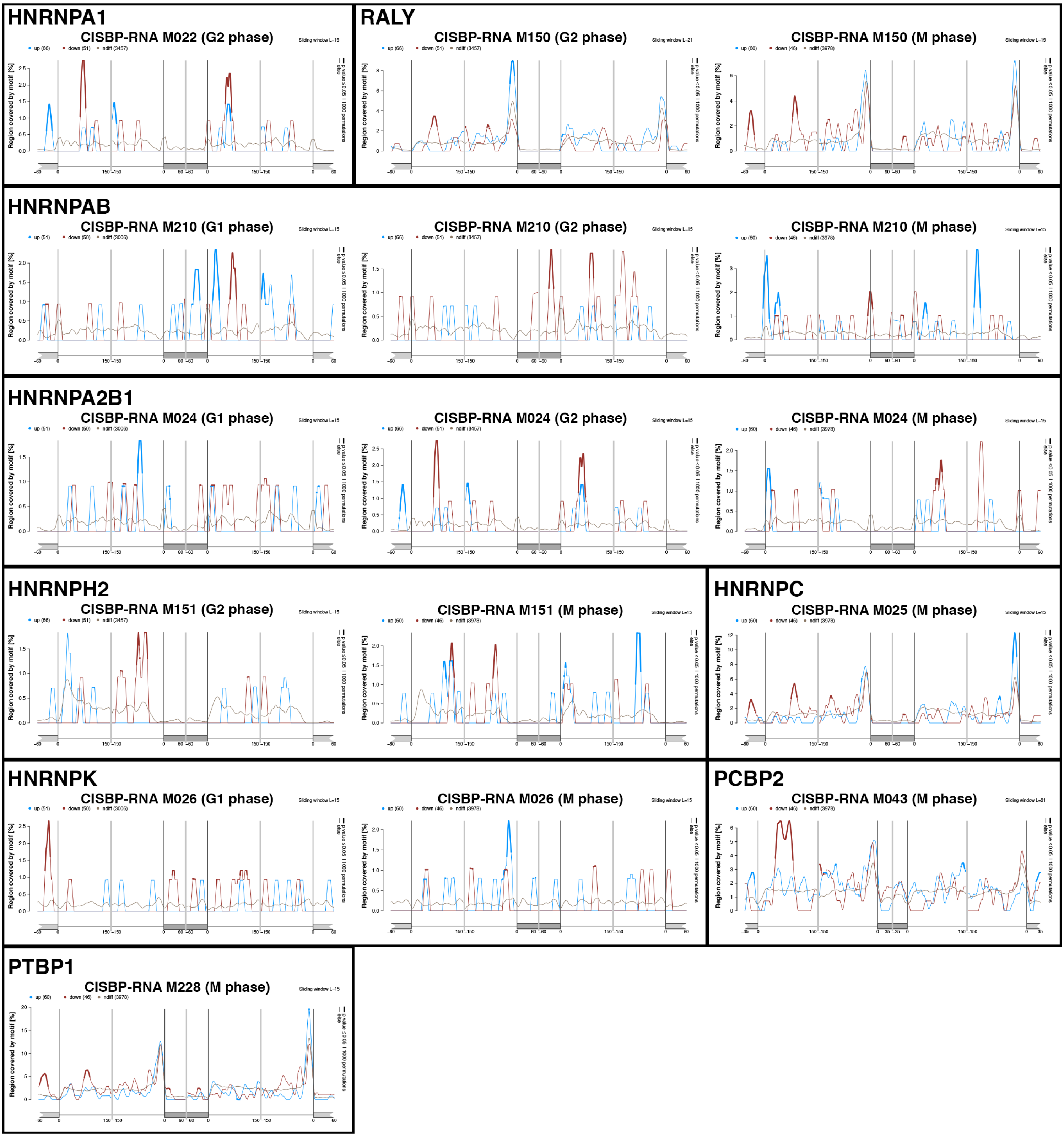

### Supplementary Figure-S8

**A**

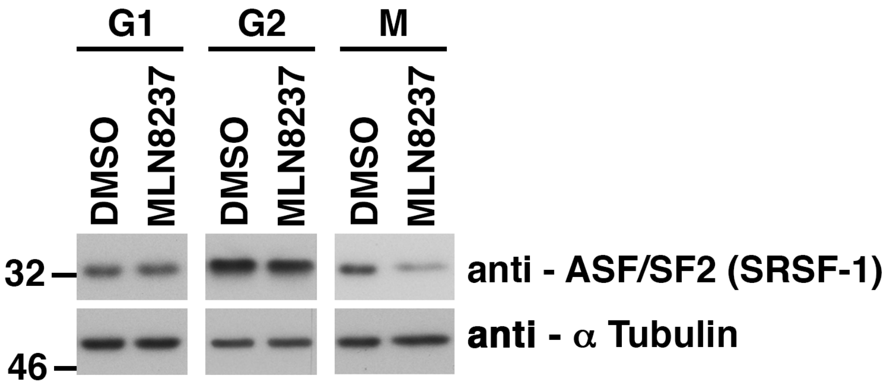
